# Interfacial tension and growth both contribute to mechanical cell competition

**DOI:** 10.1101/2025.01.09.632135

**Authors:** Léo Valon, Alexis Matamoro-Vidal, Alexis Villars, Romain Levayer

## Abstract

Tissue plasticity and homeostasis rely on the mutual interplay between cell behaviour and mechanical inputs^1^. Yet, mechanical stress can also contribute to the evolution of some pathologies, notably by accelerating pretumoural cell expansion through the process of mechanical cell competition^2–5^. Mechanical cell competition is a conserved process in which one cell population is preferentially eliminated when mixed with another cell population due to its higher sensitivity to mechanical stress^2,3,5–8^. Most of the recent theoretical and experimental explorations of mechanical cell competition focused so far on the contribution of growth and pressure to cell elimination and were limited to few genetic contexts, including the activation of Ras *in vivo*^2,8^, and the mutation of the polarity gene *scribble* in mammalian cell culture^3,4,9^. However, it remains unclear whether other oncogenes can trigger similar mechanisms and whether growth is generally the only central regulator of cell compaction and cell elimination. Using the *Drosophila* pupal notum (a single layer epithelium), quantitative live imaging and vertex modelling, we revisited the mechanisms contributing to cell compaction and cell elimination during mechanical cell competition. Doing so, we outlined the co-existence of two modes of wild type (WT) cell compaction near oncogenic cells, namely the compaction driven by growth versus local compaction driven by increased tension at tumoural/WT cell interfaces in zones of high curvature (similar to “Laplace pressure”). We highlighted distinctive signatures in cell deformation and cell elimination distribution that can delineate these two modes of compaction, and we recapitulated them *in silico* and *in vivo* using genetic backgrounds affecting growth and/or interfacial tension independently. Altogether, this study reveals for the first time the contribution of interfacial tension-driven compaction to mechanical cell competition and outlines the co-existence of various modes of compaction during cell elimination and pretumoural clone expansion.

## Results

Previously, we outlined the process of mechanical cell competition in the pupal notum using the conditional activation of an active form of the oncogene Ras (Ras^V12^) in clones, and revealed local compaction and elimination of WT cells several cell rows away from the clones^2,8^. We decided to characterise more extensively the features associated with WT cell elimination in proximity of Ras^V12^ clones. As previously reported, activation of Ras over 12 hours is sufficient to observe clone expansion (**Fig. 1 C-F**, **Fig. S1 A-C**, **movie S1**) leading to an averaged area expansion of roughly 50% over 12h. This correlated with compaction and increased death rate in the neighbouring WT cells from 1 to 3 cell rows away from clone border (**Fig. 1A-F**, **movie S1,** 30-40% area reduction, up to 3 folds increased death rate). By increasing the proportion of tissue coverage by Ras^V12^ cells, we also noticed a clear modification of the shape of WT cell patches surrounded by Ras^V12^ cell over time, which tends to round up (**Fig. 1G,H, movie S2**, significant increase of patches circularity over 10 hours). This was indicative of a cell sorting phenomenon which can be driven by the modulation of interfacial tension between mutant and WT cells^10,11^ (**Fig. 1I**). Accordingly, we observed a distinctive increase of line tension at clone interface using force inference^12^ (**Fig. S1 D-G**) and a clear enrichment of non-muscle MyosinII light chain (MyoII) at the interface between WT and Ras^V12^ cells (**Fig. S1 H-J**). This is in good agreement with previously published junction tension estimation by laser ablation the pupal notum^11^ and the observed accumulation of actomyosin at Ras clone interfaces in the wing imaginal discs^13,14^. At the interface between two fluids, interfacial tension can generate a difference in pressure called Laplace pressure which will be proportional to the interfacial tension and the local curvature of the interface (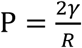, where R is the curvature radius and γ the line tension)^10,13^. By analogy, we expected WT cell patches near curved and convex interfaces to experience higher compaction relative to less curved boundaries. In agreement with this, we observed a more significant compaction of WT cell patches located in highly curved and convex area near Ras^V12^ clones relative to WT cells near flatter interfaces (**Fig. 1J-L**, **movie S3**). Focusing on WT patches with an initial elliptic shape (composed of 25-50 cells), we also observed a 2.5 fold increase of the rate of cell elimination at the highly curved tips of the WT patches relative to the central/flat areas (**Fig. 1M**, **movie S3**). Altogether, this suggested that interfacial tension combined with boundary curvature could also contribute to WT cell compaction and cell elimination near Ras^V12^ clones.

**Figure 1:**
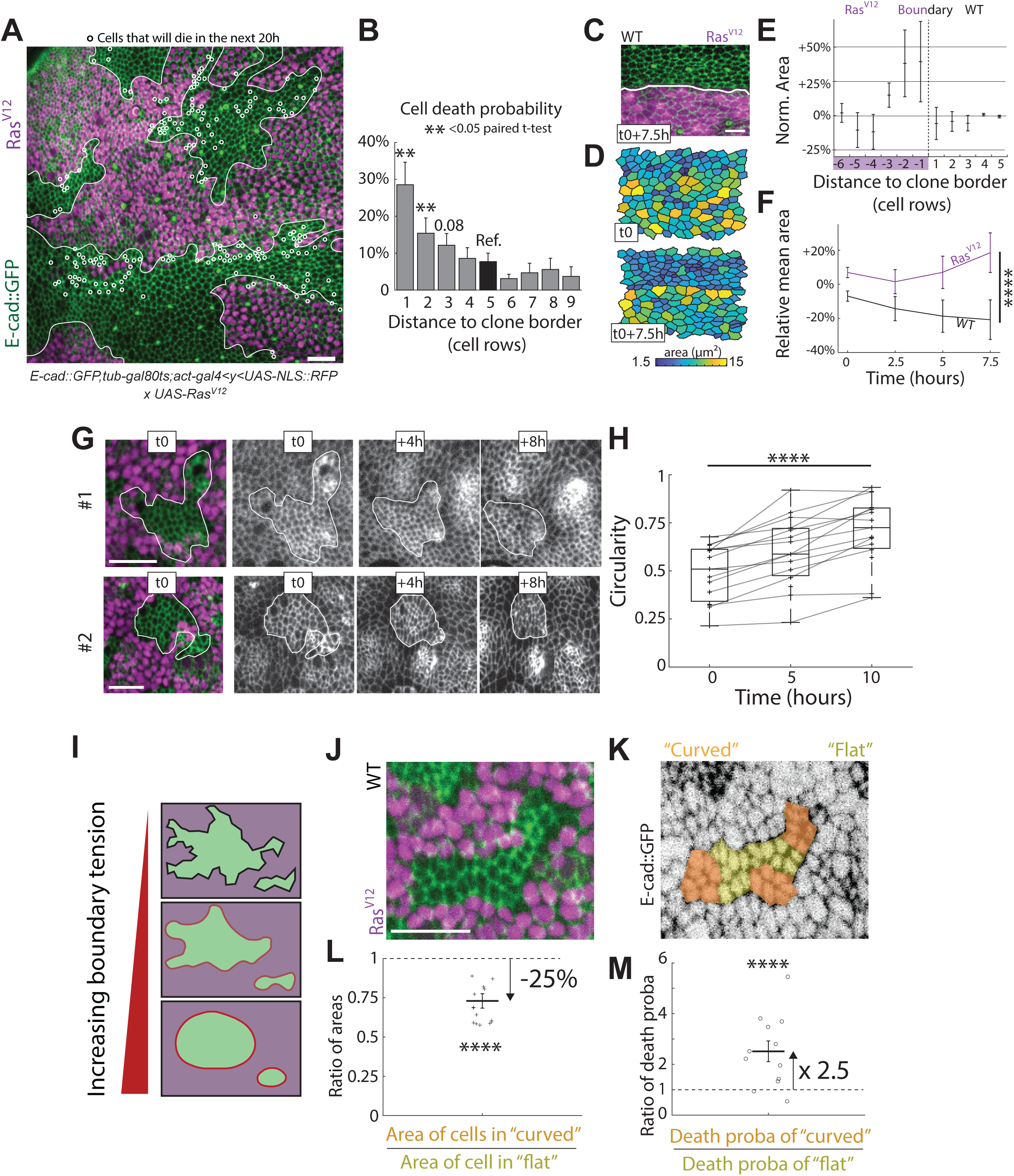
Sorting and preferential cell compaction and elimination in curved area near active Ras clones (associated with Figure S1). **A-M:** Mechanical cell competition scenario: all cells express endo-Ecad::GFP (in green) and clones (Gal4 positive, in magenta) expresses conditionally UAS-Ras^V12^. Movies start about 12 to 20 APF and last for 16 to 20h long. **A:** Clone border is underlined by white lines. Cells that will die during the 20 hours movies are highlighted by white round markers. Scale bar is 20μm. **B:** Quantification of cell death probability in WT cells as a function of distance (in cell row) to the clone boundary. Cell death numbers for each raw is divided by the number of cells present in each row at time 0 (N=626 deaths localized for over 4 nota, 4658 cells tracked). **, p<0.05 from paired t-test, error bars are s.e.m. **C-F:** Quantification of single cell compaction over time. **C:** Zoom on one clone boundary at time 0. Scale bar is 20μm. **D:** Cell apical area at 0 and 7.5 hours in Ras^V12^ and nearby WT cells of the boundary shown in **C**. Colour code is shown on the left. **E:** Normalised and averaged cell apical area about 20 to 25 hours after beginning of Ras induction as a function of distance (in cells) to clone boundary, area are normalised by mean cell area of WT cells at 4 and 5 rows away (N=393 cells, from 3 snapshots). Error bars are s.e.m. **F:** Evolution in time of the mean apical area on the 2 first rows of cells on both side of the Ras^V12^ clone boundary. N=90 cells over 4 time points from 3 movies. ****, t-test, p-value <10^-8^. Error bars are s.e.m. **G-H:** Evolution of WT cell patches shape over time. **G:** Two representative examples of WT patches surrounded by Ras^V12^ cells, clone boundaries are highlighted by a white line. Scale bar is 20μm. **H:** Boxplot of WT cell patches circularity 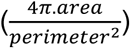 over time for 15 groups (black crosses) of WT cells from 4 experiments over 10 hours. The lines connect values for the same WT patch over time. ****, paired t-test, p-value <10^-6^. **I:** Schematics of mechanical effect of increased clone boundary tension on boundary and cell patches shape. The boundaries smoothen progressively and the patches round-up. **J-M:** Differential WT cell behaviour as a function of their localisation in curved/convex versus flat areas of the clone interface. **J:** Representative example of an elongated patch of 25 to 50 WT cells used for the quantification. Scale bar is 20μm. **K:** WT cells classification in either “curved”, in orange, or “flat” regions, in yellow. **L:** Averaged ratio of single cell areas between cells in curved versus flat regions. Single points are ratio of mean cell apical areas for a single WT patch. N=560 cells from 15 crops obtained from 4 nota. ****; t-test for the ratio being different from 1: p<10^-4^. Error bar is s.e.m. **M:** Ratio of cell death probabilities for WT cell in curved versus non-curved regions. N=199 deaths, from 12 crops obtained from 4 nota. ****, t-test for the ratio being different from 1: p=0.0035. Error bar is s.e.m.

We therefore decided to test theoretically and experimentally how interfacial tension could contribute to cell compaction and elimination relative to compaction driven by differential growth. We used a 2D vertex model^15^ (**Fig. 2A**, and **Star methods**) to explore how variation of tension at clone interface on one hand and growth on the other hand impact cell shape and cell elimination. We first confirmed that higher interfacial tension was sufficient to induce cell compaction in circular clones, and that the degree of compaction is inversely proportional to the initial radius of the clone (in agreement with the Laplace pressure law) (**Fig. S2 A,B**). We then explored the impact of the interface shape on cell compaction and cell shape evolution. When we started from an elliptic patch of WT cells (**Fig. 2A**), increasing interfacial tension was sufficient to induce significant cell area reduction specifically at ellipse tips where curvature is higher, as well as a progressive increase of the patch circularity (**Fig. 2 B-C**, **Fig. S2C**, **movie S4**) in agreement qualitatively with WT cells near Ras clones (**Fig. 1G,H**, **Fig1. J-L**, **movie S2, movie S3**). Cell compaction was also more significant in the first row of WT cells (**Fig. S2D**). Interestingly, this also led to distinctive cell shape changes, including an elongation of cells orthogonal to the interface on both side of the patch interface (**Fig. 2B**), a stretching of external cells orthogonal to the ellipse tips and a global alignment of WT cell orthogonal to the patch long axis (**Fig. 2D**, **Fig. S2C**, **movie S4).** We then explored the impact of differential growth on cell compaction, elongation and patch shape. We modulated the resting area of the surrounding cells (“mutant cells” in **Fig. 2**) to impose a progressive increase of their apical area. This led to a rapid and relatively homogenous compaction of the central elliptic patch (**Fig. 2G,H**, **Fig. S2C**, **movie S5**). Moreover, the WT cell patches do not become more circular and rather become more elongated due to compaction orthogonal to the patch long axis (**Fig. 2G,H**, **Fig. S2C**, **movie S5**), in agreement with the global elongation of cells in the direction of the patch long axis (**Fig. 2I**, **movie S5**), and in stark contradiction with what is observed upon increased of interfacial tension (clone rounding and cell elongation orthogonal to patch long axis, **Fig. 2B,D**). Interestingly, in this condition cell compaction is fairly homogeneous throughout the WT population (similar compaction in each cell row, **Fig. S2F**) and does not correlate with local clone border curvature (**Fig. 2 H**, **movie S5**), contrasting again with the deformations observed upon increased interfacial tension. Next, we introduced cell elimination in the model. Cell eliminations (T2 transitions) were triggered below a threshold area strain with a given probability in order to disperse them in time and space (**Star methods**). Doing so, we confirmed that increased interfacial tension drives the progressive elimination of cells at the tips of the ellipses, showing higher death probability in curved regions relative to less curved regions irrespective of the area strain death threshold selected (**Fig. 2E,F, movie S6**), and more death in the first row of WT cells (**Fig. S2E**), all in agreement with the experimental observations near Ras^V12^ clones (**Fig. 1M, movie S6**). On the contrary, growth-induced compaction led to a more homogeneous distribution of cell death in the patches without preferential enrichment at the patch tips (**Fig. 2J,K, movie S6**) and more elimination in the two first rows of cells (**Fig. S2G, movie S6**). Both conditions could eventually lead to a full elimination and/or fragmentation of the WT patches depending on the growth/tension characteristics and on the area strain death threshold (**Figure S2H,I**).

**Figure 2:**
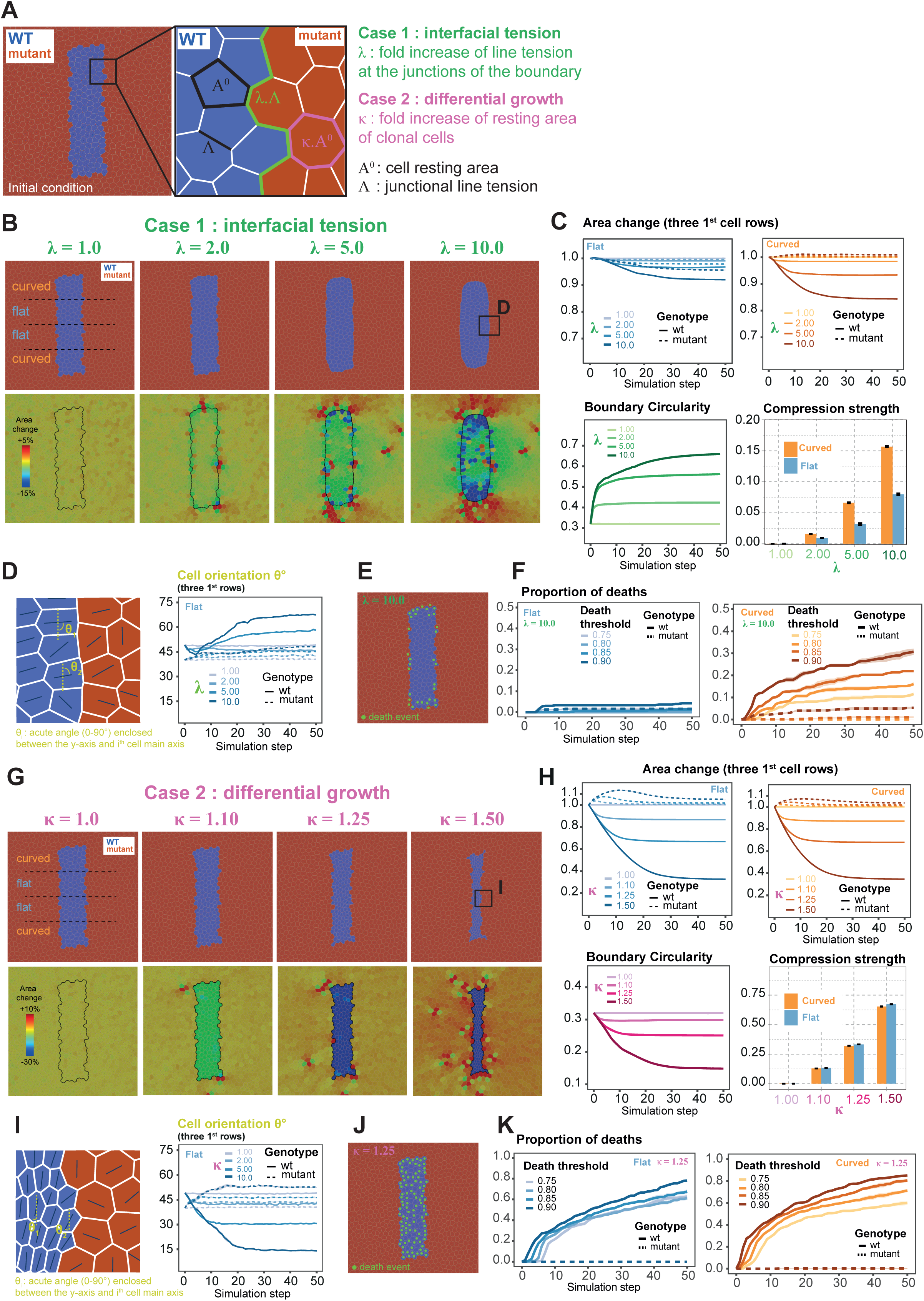
vertex modelling of mechanical cell competition (associated with Figure S2). **A:** Topology used for initial conditions of the simulations. The tissue is made of ∼3600 cells with fixed boundary conditions. In the centre of the tissue, a group of ∼150 cells is set as WT (blue cells) whereas all the other cells of the tissue are set as mutant (red cells). The inset shows the parameters used in the vertex model to simulate two different mechanical cell competition scenarios. For the “interfacial tension” scenario (Case 1) the line tension parameter of the junctions at the boundary (green line) between WT and mutant cells is increased by a factor λ with respect to the line tension Λ of all the other junctions of the tissue. For the “differential growth” scenario (Case 2), the resting area of the mutant cells is increased of a factor κ with respect to the resting area A^0^ of all the other cells of the tissue. **B-K:** Simulations outcomes from two mechanical cell competition scenarios: interfacial tension (Case 1, panels B to F) and differential growth (Case 2, panels G to K). **B:** Final state for simulations of Case 1 obtained with four different values of λ. Top panels show the topology of the tissue, and bottom panels show single cell area fold change at the final simulation step, relative to area at the beginning of the simulation. Black line is the boundary between WT and mutant cells. **C:** Top panels: cell area fold change relative to initial area (averaged over all the cells of the three first rows of cells from the boundary between WT cells and mutant cells) as a function of simulation step, λ, genotype and curvature of the boundary (flat vs. curved). See top left panel in B for a schematic of flat vs. curved regions. Bottom left panel: circularity 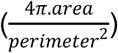 of the boundary of the WT cells population, as a function of simulation step and of λ. Bottom right panel: cell compression strength 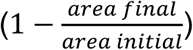 for the WT cell population as a function of λ and of boundary curvature. **D:** Left: inset from panel B (top right) showing for each cell the direction of its longest axis (blue lines, given by the direction from cell’s centroid to its farthest vertex). Cell orientation is measured by θ, the acute angle enclosed between the y-axis of the tissue plane and the direction of cell’s longest axis. Right panel: θ averaged over the three first rows of cells from the boundary as a function of simulation step, λ and cell’s genotype (WT or mutant). Data shown for the flat region only. **E:** Back tracking of dying cells. Cells that will die during the simulation (Case 1, λ = 10.0) are highlighted on the topology at simulation step 0 (green dots). **F:** Proportion of deaths for the three first rows of cells from the boundary, as a function of simulation step, genotype, curvature of the boundary and death area threshold. Data are shown for Case 1, λ = 10.0. Proportion is given relative to the number of cells of a given genotype and curvature region at simulation step 0. **G:** Final state for simulations of Case 2 obtained with four different values of κ. Top panels show the topology of the tissue, and the bottom panels show single cell area fold change at the final simulation step relative to area at the beginning of the simulation. Black line is the boundary between WT and mutant cells. Note that the colour scale of the heat-map is different from the one of panel B. **H:** Top panels: cell area fold change relative to initial area (averaged over all the cells of the three first rows of cells from the boundary between WT cells and mutant cells) as a function of simulation step, κ, genotype, and curvature of the boundary (flat vs. curved). See top left panel in G for a schematic of flat vs. curved regions. Bottom left panel: circularity 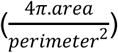 of the boundary of the WT cells population, as a function of simulation step and of κ. Bottom right panel: cell compression strength 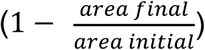 for the WT cell population as a function of κ and of boundary curvature. **I:** Left: inset from panel G (top right) showing for each cell the direction of its longest axis (blue lines, given by the direction from cell’s centroid to its farthest vertex). Cell orientation is measured by θ, the acute angle enclosed between the y-axis of the tissue plane and the direction of cell’s longest axis. Right panel: θ for the three first rows of cells from the boundary as a function of simulation step, λ and cell’s genotype (WT or mutant). Data shown for the flat region only. **J:** Back tracking of dying cells. Cells that will die during the simulation (Case 2, κ = 1.25) are highlighted on the topology at simulation step 0 (green dots). **K:** Proportion of deaths for the three first rows of cells from the boundary, as a function of simulation step, genotype, curvature of the boundary and death area threshold. Data are shown for Case 2, κ = 1.25. Proportion is given relative to the number of cells of a given genotype and curvature region at simulation step 0. All quantifications are averages +/− sem obtained by running 5 times the simulations using different seeds in the Monte Carlo random walk.

Altogether, the simulations suggested that high interfacial tension and differential growth are both sufficient to compact and eliminate cells, while exhibiting distinctive features. Increased interfacial tension induced a higher cell compaction and cell elimination in curved regions, a preferential elongation of cells orthogonal to the patch long axis, and a progressive rounding-up of the cell patch. In contrast, differential growth led to a more homogenous and isotropic compaction and elimination, cell elongation parallel to the patch long axis, and the absence of patch rounding.

To validate these predictions, we tried to find experimental conditions that could impact growth and line tension independently (unlike Ras^V12^ activation which modulates both). Activation of Yki (the fly orthologue of Yap) can promote growth as well as cell survival^16^. Moreover, Yki activation in clones was previously associated with neighbouring cell elimination both in *Drosophila* wing discs^17^ and in the pupal notum^18^. Conditional expression of a constitutively active form of Yki (Yki^CA^, Yki^S111A,S168A,S250^ triple mutants^19^) in large clones led to a rapid expansion of clone surface and compaction of the neighbouring cells (**Fig. 3 A-D,H,I Figure S3A**, **movie S7**), and high death rate in WT cells up to 4 cell rows away from the clone (**Fig. 3 E-G**, **movie S7**). Contrary to competition driven by Ras^V12^, we did not observe a smoothening of the WT/clone interface nor the rounding-up of WT patches (**Fig. 3 H-J, movie S8**) suggesting that compaction is here shape conservative and not associated with sorting. Accordingly, we could not find any significant increase of line tension at clone interface using force inference and MyoII distribution (**Fig. S3 B-D**). Focusing on elliptic patches of WT cells, we found that the compaction was associated with a progressive elongation of WT cells along the long axis of WT patches, in agreement with the predictions of the vertex model with differential growth (**Fig. 3. K,L**, **movie S8**, compare with **Fig. 2G,I**, **movie S5**). More importantly, in contrast to Ras^V12^ competition and the model of high interfacial tension, we could not find any difference for WT cell compaction and death probability in curved versus flat interfaces near Yki^CA^ clones (**Fig. 3 M-P, movie S9**). The more homogeneous distribution of cell death in WT patches fits the predictions of the model based on differential growth (**Fig. 2H,J,K**, **Fig. S2 F,G, movie S5 and S6**). Altogether, we found that WT cell compaction and elimination induced by Yki^CA^ clones fit all the features observed in the vertex simulations of growth-induced compaction (**Fig. 2G-K**).

**Figure 3:**
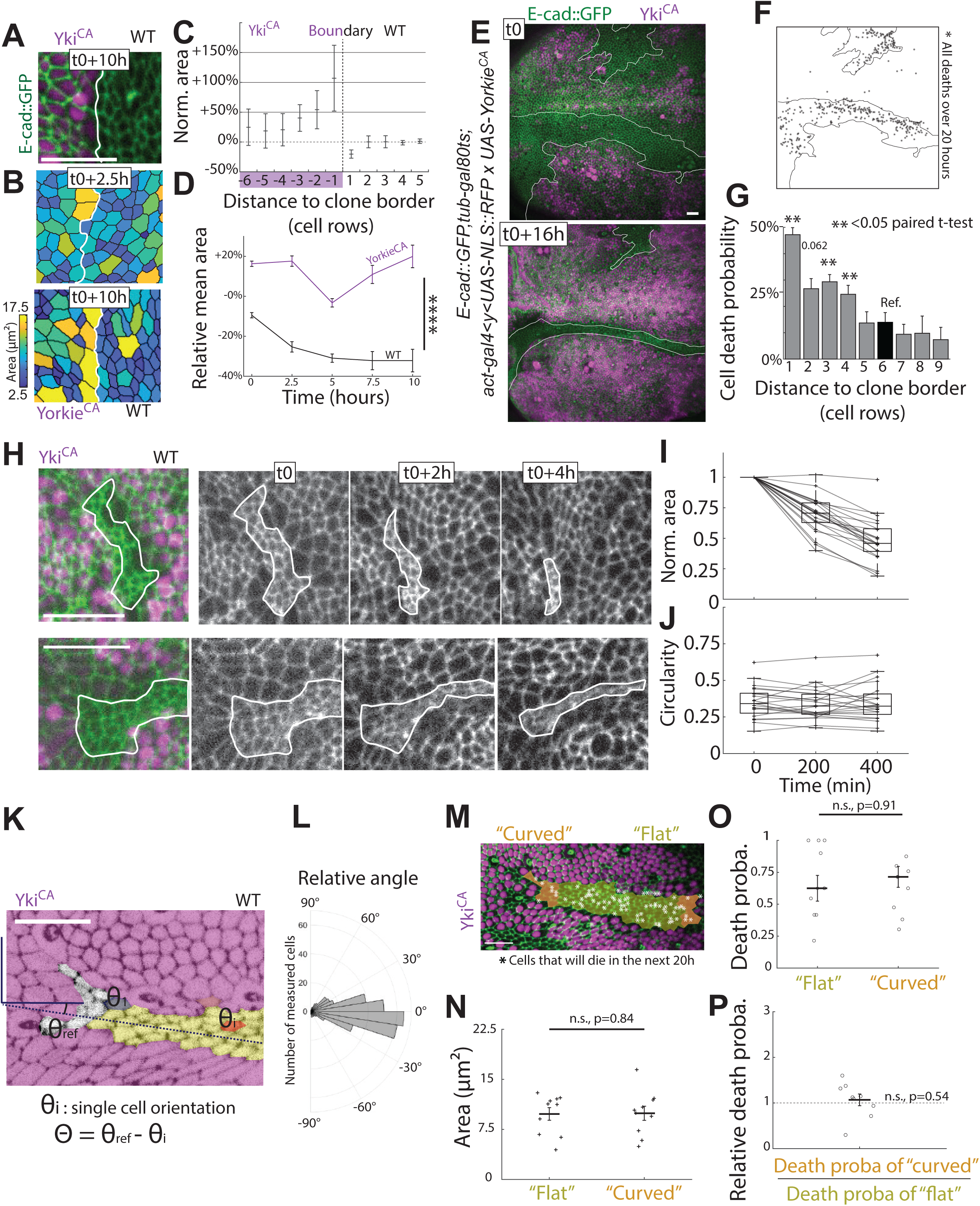
Yki clonal activation and growth-driven mechanical competition (associated with Figure S3 and S4) **A-P:** Differential growth generated by the overexpression of an active form of Yorkie (Yki^CA^). All cells express E-cad::GFP (green). Magenta: conditional overexpression of UAS-Yki^CA^, expression started 8h before the start of imaging phase which last 16-20h. **A-D:** Quantification of single cell compaction over time. **A:** Zoom on one boundary between Yki^CA^ and WT population 10h after Yki induction. Scale bar is 20μm. **B:** Segmentation of two snapshots of a Yki^CA^ clone interface separated by 10 hours, colour code (on the left) shows the apical area. **C:** Quantification of cell apical areas as a function of distance (in cells) to clonal boundary between Yki^CA^ clone and WT population 20 to 25 hours after Yki induction normalised by mean cell area of WT cells at 4 and 5 cell distance (N=440 cells, 3 nota). **D:** Quantification of the evolution of mean apical area of the 2 first cell rows at both side of the boundary. N=90 cells, 5 time points from 3 nota. Time points are separated by 2.5 hours. ****, t-test, p-value <10^-16^. Error bars are s.e.m. **E:** Snapshots of a nota at time t0 (top) and t0+16 hours (bottom) expressing E-cad::GFP and activating Yki^CA^ in magenta clones. Approximative clone border is highlighted by white lines. Scale bar is 20 μm. **F:** Representation of clone boundaries. Black asterisk: back tracking of the localisation of all the cells that will die during the 20 hours movie. **G:** Quantification of cell death probability as a function of distance to the border (in number of cells). Cell death numbers in each row is divided to the initial number of cells at the same distances (N=1074 deaths localized for over 4 nota normalised by 4463 cells). **, p<0.05 from paired t-test with cell death probability at row 6 used as reference. **H-J** Evolution of WT population shapes over time. **H:** Snapshots of two representative examples with border of WT population surrounded by Yk^iCA^ clones (magenta) highlighted by a white line. Images are separated by 2 hours. E-cad is shown in green and gray. Scale bar is 20μm. **I-J:** Boxplot of WT cell apical area (**I**) (normalised by the first time point) and circularity (**J**) 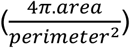 for 22 groups of WT cells surrounded by Yki^CA^ cells (single curves) from 6 experiments over about 6 hours. The lines connect the value coming from the same group of cells. **K-P:** Quantification of single cell shape, compaction and death as a function of curvature in the WT cell groups. **K:** Schematic of the determination of single cell orientation with reference the main orientation of the WT group. Cells at the tips are excluded from the analysis. WT group main orientation is represented by a dashed line. Scale bar is 20 μm. **L:** Polar distribution of the orientation of WT cells surrounded by Yki^CA^ cells 20 to 25 hours after Yki^CA^ induction. N=342 cells from 16 groups from 6 nota. **M:** Example of an elongated WT patch composed of ∼25-50 cells used for the curvature analysis. Representation of WT cells classification in either “curved” (orange) or “flat” (yellow) area. White stars show dying cells. **N:** Mean apical area of cells in “flat” and “curved” areas. One dot=one WT patch. Paired t-test: n.s., not significant, p=0.84. **O-P:** Cell death probability of WT cells in “flat” or “curved” areas (**O**) and ratio of cell. N=219 cell deaths, from 9 groups from 4 nota. Error bars are s.e.m. **O:** Mean of death probabilities. Paired t-test: n.s., not significant, p=0.91. **P:** Determination of ratio of cell death probabilities. t-test for testing difference to 1: n.s., not significant, p=0.54.

We then examined conditions affecting interfacial tension independently of growth. Previously, ectopic expression of N-cadherin (N-cad) was shown to stimulate MyoII accumulation and increased tension at the border with cells that do not express N-cad^20^. We recapitulated these observations in the pupal notum using conditional expression of N-cad in clones: clone borders are progressively smoothening and are associated with an upregulation of tension and MyoII recruitment at the boundary (**Fig. 4 A-F**, **movie S10**, **Figure S3E-G**), hence leading to progressive clone/WT patches rounding and smoothening of finger-like interfaces (**Fig. 4C,D,F, movie S10**). Strikingly, N-cad induction led to rapid compaction of both WT and N-cad cells located near convex borders (**Fig. 4G-I**, **movie S11**), which was also associated with cell elimination on both sides (**Fig. 4 A,C**, **movie S11**). Accordingly, cells were roughly 30% smaller (**Fig. 4I**) and death probability was twice higher (**Fig. 4 J,K**) near convex boundary compared to region near flat boundary, in agreement with the predictions of the vertex model with high-interfacial tension (**Fig. 2B,C,F**). Similarly, N-cad induction recapitulated the elongation of first rows of cells orthogonal to the clone borders (**Fig. 4L,M** and **Fig. 2D**). Surprisingly, we observed two important features that were not predicted by the vertex model: a relatively higher cell apical area of the first row of WT and N-cad cells near the interface (**Fig. 4G,H,L**), and a lower death rate in the first row of WT and N-cad cells (**Fig. 4B**), while the model predicted that the first row of cells were the most compacted and the more susceptible to die near convex regions (**Fig. S2D,E**). We therefore tried to find which features might be missing in our initial vertex simulation of increased interfacial tension. A closer look at MyoII distribution as well as force inference not only revealed increased tension at clone boundary, but also an unexpected decrease of MyoII and tension levels in junctions orthogonal to clone boundary (**Fig. S3E-G**). We therefore implemented a similar reduction of tension in junctions orthogonal to clone boundary combined with the increase of clone boundary tension in our vertex model. This was sufficient to recapitulate the relative increase of cell area at the boundary and the reduction of cell elimination in the first cells rows near convex boundary (**Fig. S4**, **movie S12**), suggesting that the pattern of cell death could be fully explained by the combination of high interfacial tension and tension reduction in orthogonal junctions. Altogether, this provides to our knowledge the first clear experimental evidence that increased line tension is sufficient to drive cell compaction and cell elimination *in vivo*.

**Figure 4:**
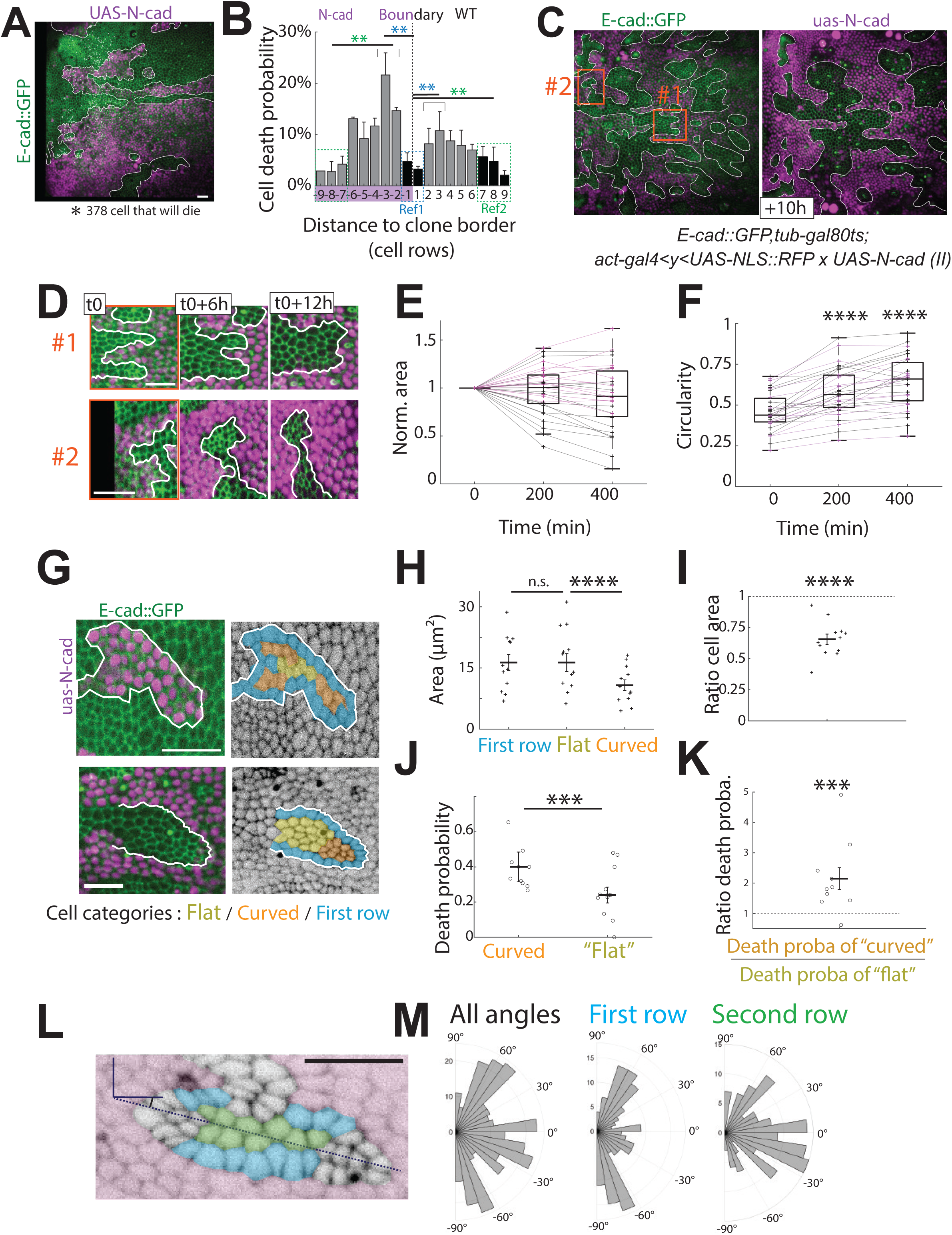
N-cad overexpression drives sorting and curvature dependant cell elimination (associated with Figure S3 and S4) **A-M:** Mechanical cell competition scenario generated by differential junctional tension via the overexpression of N-cadherin. All cells express E-cad::GFP (in green) and N-Cadherin overexpressing cells are in magenta. N-cadherin expression started 8h before the start of imaging phase that lasts 16-20h. **A-B:** Localisation of cells that die as a function of distance to the boundary between populations. **A:** Snapshot of a z-local projected movie. White line highlights the clone boundaries. White asterisks: back tracking of the localisation of the 378 cells that will die during the next 16 hours. Scale bar is 20μm. **B:** Quantification of cell death probability as a function of distance to the boundary (in number of cells). Negative distance values are for cell rows inside the N-cad clones. Cell death numbers in each row is divided by the initial number of cells in this row (N=644 deaths localised for over 4 nota normalised by 7893 cells). Two rounds of t-test for death probability of rows (−3, −2) and (+2, +3) have been performed against reference Ref1 and Ref2 rows. **, p<0.05. **C-D:** Evolution of clone shape boundaries for two images separated by 10h. Clone boundaries are highlighted by white lines. **D:** Two representative examples (red square in **C**) with border of WT population represented by a white line. Images are separated by 6 hours. Scale bar is 20μm. **E,F:** Analysis of N-cad clone (magenta curves) and WT (black curves) groups boundary shape evolution over time. **E:** Normalised area. **F:** circularity, ****, p< 10^-5^, paired t-test (N=31 groups of cells from 8 nota). Single lines represent the evolution of a clone/group of cell. Box plots show the median, the 25% and 75% percentile and min and max values. **G-K:** Quantification of single cell compaction and probability of cell death as function of boundary curvature. **G:** Representative examples. Clone boundaries are represented by white lines. Cells are classified among 3 independent categories: in the first cellular row (in blue), in a region close to a curved boundary (in orange), in a region close to a “flat” boundary (in yellow) in N-cad clones (top) or WT patches surrounded by N-cad cells (bottom). Scale bars are 20 μm. **H:** Quantification of mean cell area for cells in the 3 different categories after 20 to 25 hours of N-cad induction. Paired t-test, n.s., not significant, p=0.56; ****, p<0.0005 (N=140, 125 and 157 cells, from cell 12 groups from 6 nota). **I:** Quantification of the ratio of mean cell areas for “curved” over “flat” category. ****, p<10^-5^, paired t-test against “1”, one cross = one group of cells. **J:** Quantification of cell death probability for cells in the 2 categories “flat” and “curved”. **, p=0.0056, paired t-test (N=96 cell deaths events from 11 groups of cells from 6 nota), one dot = one group of cells. **K:** Ratio of cell death probabilities for cells in “curved” over “flat” categories. ***, p=0.014, paired t-test against “1”. **L:** Example showing cell orientation in a WT cell patch surrounded by N-cad cells relative to the long axis of the patch after 20 to 25 hours of N-cad induction. The green cells (inner row) and the blue cells (first cell layer) are the one used for the analysis. We excluded the cells in the most curved regions as the angle relative to boundary orientation is more difficult to define. Scale bar is 20 μm. **M:** Polar distribution of cell orientation in the first row (blue) and second row (green) of compacted population. N=166 “first row” and 150 “second row” cells from 16 groups from 6 nota.

As previously characterised in the pupal notum^2,8^, we confirmed that compaction driven by increased growth and/or line tension triggers caspase activation (using effector caspase sensor GC3Ai^21^, **Fig S5A-D, movie S13**) and that caspase activation is required for cell extrusion (combining p35 with N-cad in clones blocked cell elimination, **Fig. S5F,G**). Ras activation in clones in larval wing disc was previously associated with JNK pathway activation at the clone interface which is responsible for WT cell elimination^22^. However we did not observe any obvious sign of JNK activation at Ras^V12^ or Yki^CA^ clones boundary and no correlation with WT cell compaction and elimination in the pupal notum (**Fig. S6A,B**). This suggests either that cell elimination is driven by a different mechanism in the notum compared to the wing disc, and/or is based on different mechanisms on the shorter timescales of our experiments (hours versus days in the larval wing disc). We also noticed that the level of sorting and the characteristic cell shape in WT patches surrounded by Ras clones vary depending on the localisation in the notum (with more pronounced sorting in posterior regions), suggesting that the relative contribution of growth versus line tension may also depend on the tissue localisation and the underlying patterning in the Ras context (**Fig. S6C**), as observed in the wing disc^23^. Interestingly, a very similar mechanism of elimination was outlined for MDCK cells depleted for the tight junction protein ZO1^24^, which are eliminated through the formation of an actomyosin purse-string and cell compaction, suggesting that mechanical competition driven by interfacial tension is conserved in mammals.

Altogether, we demonstrate experimentally and theoretically for the first time that different modes of compaction can co-exist during mechanical cell competition, including compaction and elimination driven by increased interfacial tension and compaction driven by growth. These different modes can co-exist in certain background (e.g.: upon Ras activation) and show distinctive features in term of cell shape and localisation of cell elimination (**Fig. 5**). Future work will help to reveal the relative contribution and potential synergy of these two mechanisms upon activation of other oncogenes and in vicinity of solid tumours.

**Figure 5:**
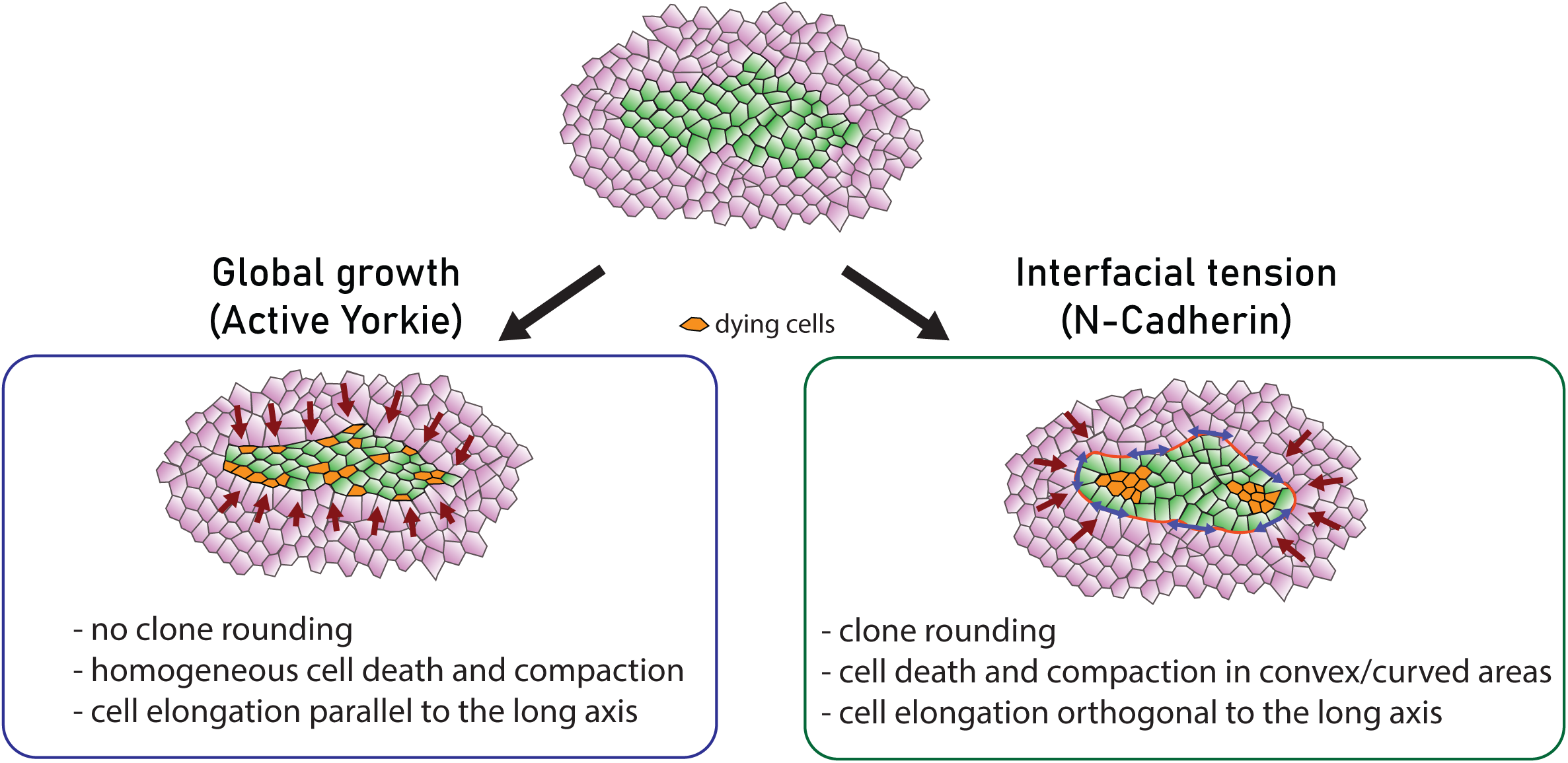
Summary of the features of the two modes of mechanical cell competition. Schematic of the two modes of compaction: compaction driven by growth (left) or by interfacial tension (right). Dying cells are shown in orange and the main cell shape and interface shape evolution features are described in the two boxes.

## Acknowledgements

We would like to thank members of the CDEH group for constructive comments on the manuscript. Work in RL lab is supported by the Institut Pasteur, the ERC starting grant CoSpaDD (Competition for Space in Development and Disease, grant number 758457), the European Union (ERC, PrApEDoC, 101085444), Views and opinions expressed are however those of the author(s) only and do not necessarily reflect those of the European Union or the European Research Council. Neither the European Union nor the granting authority can be held responsible for them, the Cercle FSER and the CNRS (UMR3738), the ANR PRC CoECECa, the ANR PRC MAPEFLU, and the ANR-10-LABX-0073. L.V. was supported by a Marie Sklodowska-Curie postdoctoral fellowship (MechDeath, 789573).

## Authors contributions

LV, AMV and RL designed the project. LV performed all the experiments and analyses, AMV performed all the vertex simulations and analyses. AV adapted the force inference method. RL supervised the project and provided funding. All the authors contributed to the article writing and editing.

## Star Methods

### Ressource availability

#### Lead contact

Further information and requests for resources and reagents should be directed to and will be fulfilled by the lead contact, Romain Levayer.

#### Material availability

All the reagents generated in this study will be shared upon request to the lead contact without any restrictions.

#### Data and Code availability

All code generated in this study and the raw data corresponding to each figure panel (including images and local projection of movies) will be accessible on a repository upon final publication of the article. Further information about the dataset can be requested to the lead contact.

### Experimental model and subject details

#### Drosophila melanogaster husbandry

All the experiments were performed with *Drosophila melanogaster* fly lines with regular husbandry technics. The fly food used contains agar agar (7.6 g/l), saccharose (53 g/l) dry yeast (48 g/l), maize flour (38.4 g/l), propionic acid (3.8 ml/l), Nipagin 10% (23.9 ml/l) all mixed in one liter of distilled water. Flies were raised in plastic vials in a room set at 22°C unless specified in the legends and in the table below (alternatively raised in incubators at 18°C or 29°C, 12/12h light cycles). Females and males were used without distinction for all the experiments. We did not determine the health/immune status of pupae, adults, embryos and larvae. Flies were not involved in previous procedures, and they were all drug and test naïve.

#### Drosophila melanogaster strains

The strains used in this study and their origin are listed in the table below.

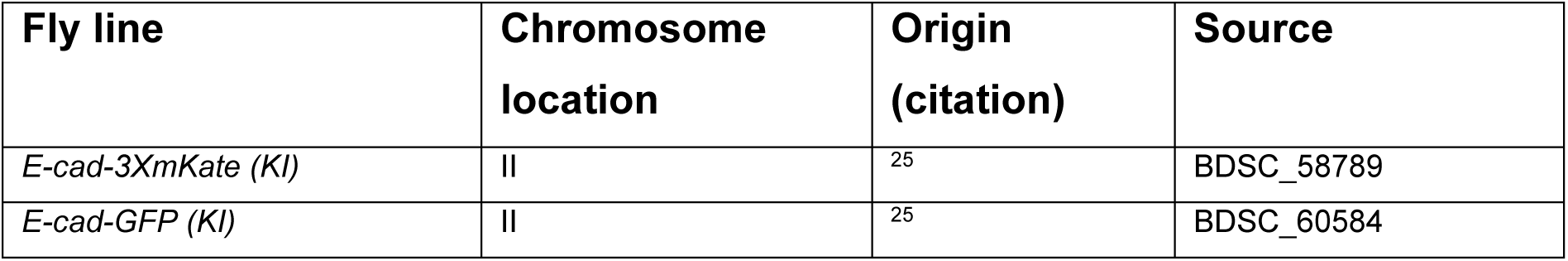
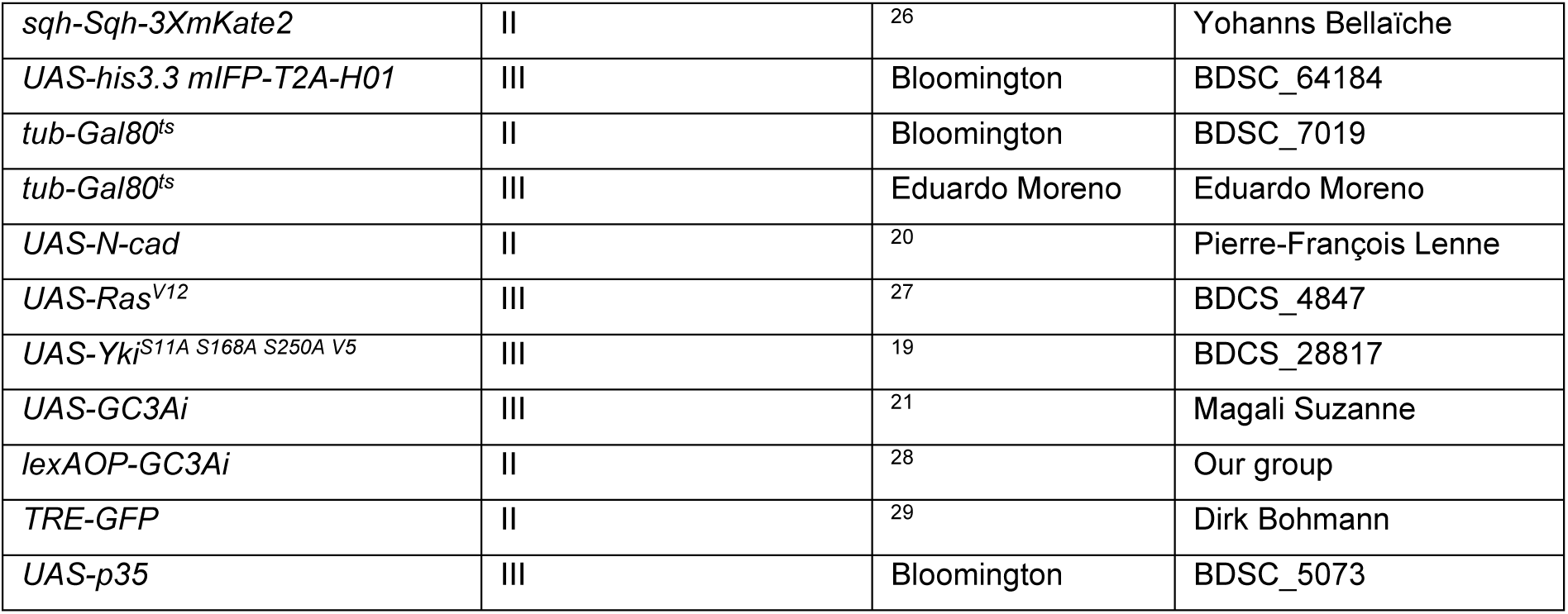

The exact genotype used for each experiment is listed in the next table. ACI: time After Clone Induction, APF: After Pupal Formation, n: number of pupae/adults.

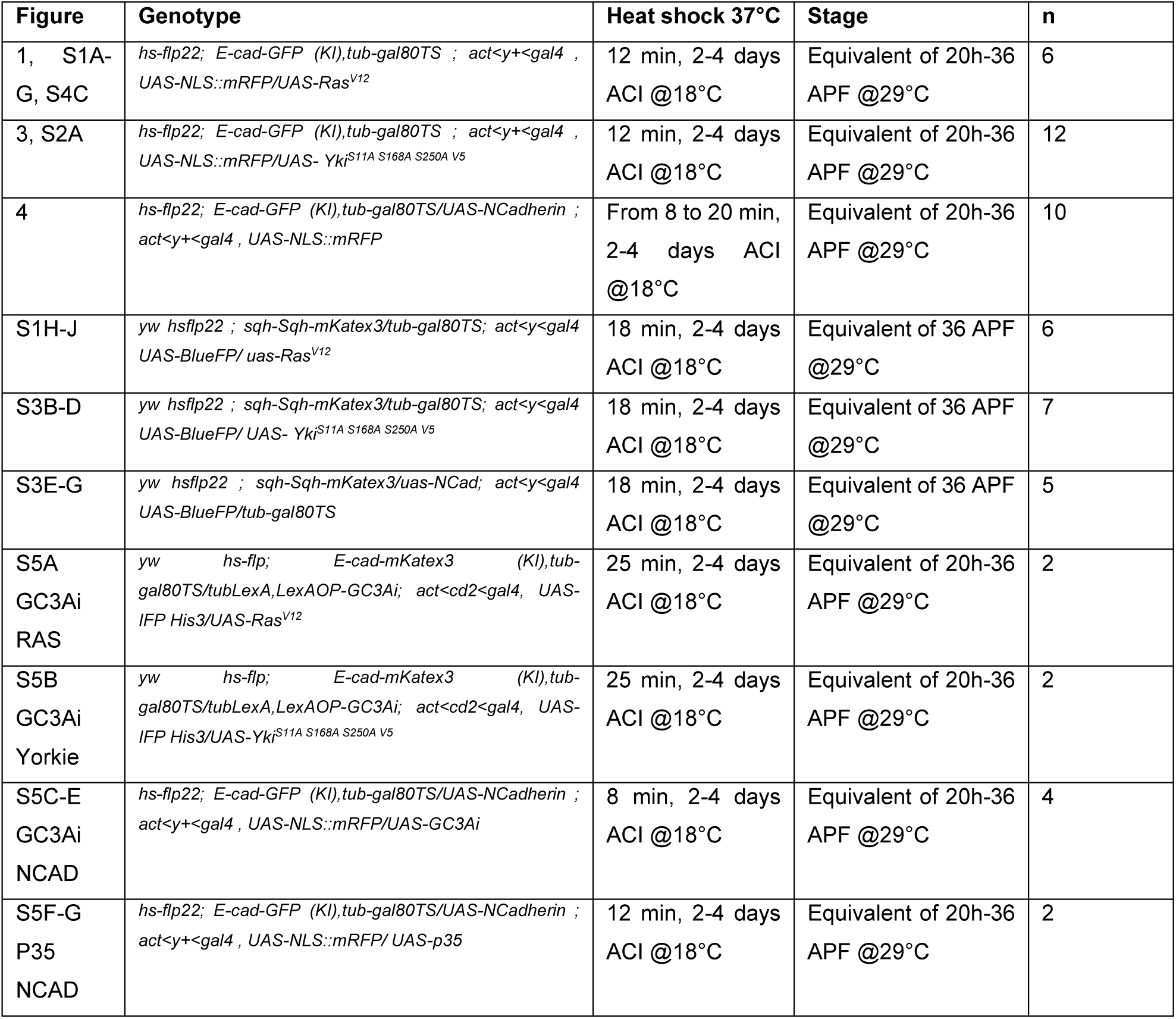
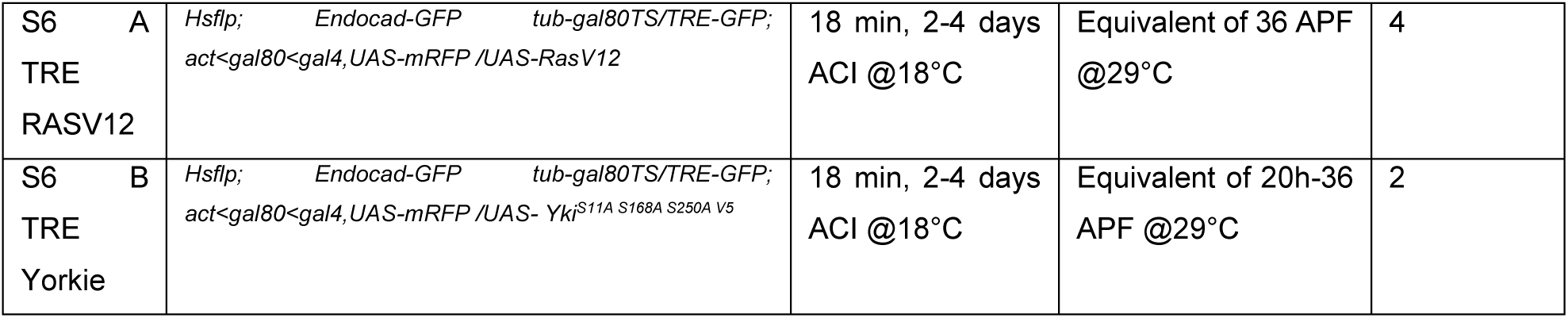

### Live imaging and movie preparation

After Heat Shock at 37°C, crosses were kept at 18°C for 2 to 4 days. The pupae were collected at the early stage (0-6 hour after pupal formation),and transferred at 29° for 6-8 hours, then glued on double sided tape on a slide and surrounded by two home-made steel spacers (thickness: 0.64 mm, width 20×20mm). The pupal case was opened up to the abdomen using forceps and mounted with a 20×40mm #1.5 coverslip where we buttered halocarbon oil 10S. The coverslip was then attached to spacers and the slide with two pieces of tape. Most timelaps movies last at least 16 hours with each positions visited every 5minutes. Pupae were imaged on a confocal spinning disc microscope (Gataca systems) with a 40X oil objective (Nikon plan fluor, N.A. 1.30). The autofocus was performed using fluorescence of cadherin as a reference using a custom made Metamorph journal. Movies were performed in the nota close to the scutellum region containing the midline and the aDC and pDC macrochaetae. For myosin quantifications and force inference measurements, a few static images were acquired with an LSM880 using an oil 40X objective (N.A. 1.3) equipped with a fast Airyscan (Zeiss).

Movies shown are adaptive local Z-projections. Briefly, E-cad plane are used as a reference to locate the plane of interest (plane of the epithelium/plane of cells) on sub windows. This procedure was either applied using a custom Matlab routine^8^ or through the Fiji plugin LocalZprojector^30^. Fluorescent signals were then obtained by projecting maximum of intensity on a few micrometers around a focal point which was centered on the surface of interest or with an offset of a few micrometer.

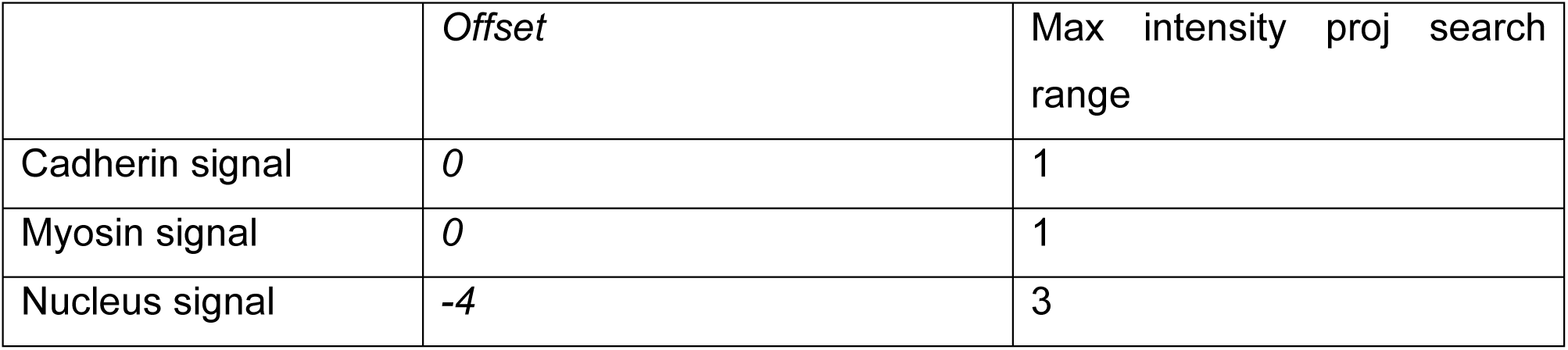
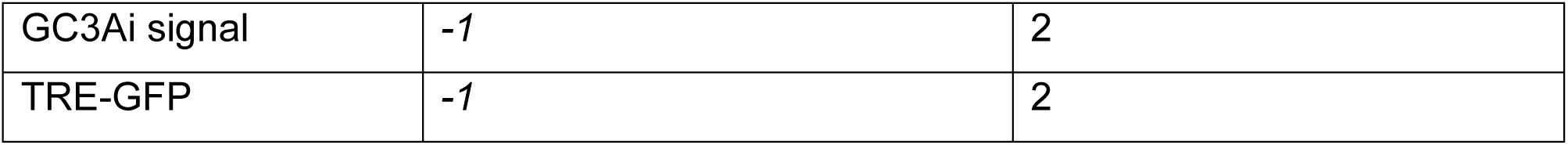

### Image analysis

#### Measurement of cell shape and compaction

To get single shape measurement (in some cases), fluorescent signal of apical cues (cadherin or myosin) were first projected, then segmented using Epyseg^31^. Segmentation were then curated by hand using the Tissue Analyzer plugin from FiJi ^32^ and then saved. Cells of interest were clicked by hand on FiJi, and list of cells were saved. Cells positions, shapes, and morphometrics were extracted and plotted using Matlab.

#### Myosin levels

From segmented myosin junctions, we also obtain boundaries of single cells and measure myosin levels at junctions with Matlab.

#### Group of cell shapes

To get “area” and “circularity” of large group of cell over time, group contour where drawn by hand on FiJi and shape characteristics were extract via Matlab. For the determination of group areas over time, groups of cells were sometimes simplified by drawing triangles in-between SOPs for WT and clonal populations.

#### Calculation of cell death probability

To determine probability of death, most (if not all) deaths were pointed by hand over the duration of the movie. Position of death were determined at the last time point the cells have an non-zero apical area. Death probabilities at a given distance were then determined by considering the number of death at that distance divided by the initial number of cell at that given distance at the first time point of the movie. Death probabilities in a zone of interest are determined by considering the number of deaths in a given zone divided by the initial number of cells in that zone.

#### Force inference

Junctional tensions and cell pressures were inferred using Bayesian inference as implemented in^12^ based on mathematical formalisms of the method developed by^33^. Tissues were segmented using TissueAnalyzer and manually corrected. Then bonds and cells files were extracted from TissueAnalyzer to csv files. MATLAB Codes from^12^ were adapted and simplified in order to load these files and track the junctions/cells of interest. Then inference was run for all segmented images.

Cells and junction of interest – at the boundary vs. in the clone vs. in the WT population – were clicked using FiJi and applied on cell pressure maps and cell junction tension maps using Matlab.

### Vertex modelling of mechanical cell competition

To infer signatures of the different processes at play during Ras^V12^ mediated mechanical cell competition, we used our vertex model program^5,34,35^ which is based on the existing framework for the study of developmental processes in the epithelial tissues of *Drosophila*^36 37^.The model was implemented in gfortran, using openGL and R to visualize the outputs.

In the vertex model, only the apical sides of the cells are considered. Cells are represented as 2D polygons, made of vertices connected by edges. The vertices can move over time as a result of intra- and inter-cellular mechanical forces. The movement of the vertices is implemented by comparing the mechanical energy of a vertex in its current position (x, y) with the energy of a randomly chosen point nearby (x+δd, y+δd) with δd ꞓ[0,0.005]. When the energy in the new position is smaller, then the movement is accepted as the new vertex location. When the energy is bigger, the movement is accepted with probability P_accept_ = 0.05 in order to introduce stochastic fluctuations.

The energy (E) of a vertex i is given by

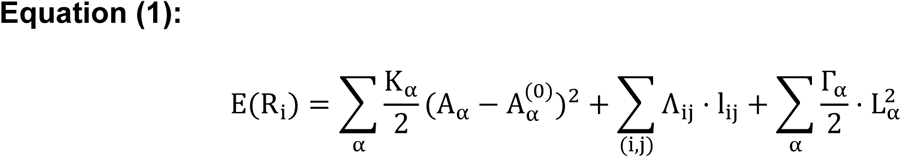

where R_i_ = (x_i_, y_i_) is the position of the vertex i. The first and the third summations are over all the cells α in which the vertex i is present, and the second summation is over all the cell edges {i, j} in which the vertex i is present. A_α_ is the apical area of the cell α and K is the area elasticity modulus, which is assumed to be equal for all the cells in our simulations. A_α_^(0)^ is the resting area of the cell α. The distance and the line tension between the pairs of vertices {i, j} are denoted l_ij_ and Λ_ij_, respectively. The third term includes the perimeter of the cell α (L_α_) and the perimeter contractility coefficient (Γ_α_). By choosing 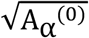 as a unit of length and (KA_α_^(0)^)^2^ as a unit of energy (as in ^36^), dividing both sides of Equation (1) by (KA_α_^(0)^)^2^ results in the following dimensionless equation:

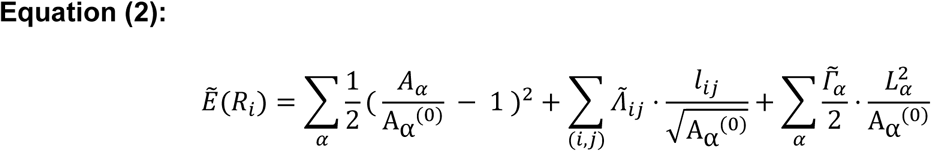

Where (A_α_⁄A_α_^(0)^), 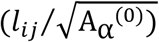 and (L_α_^2^⁄A_α_^(0)^) are, respectively, dimensionless area, bond length and perimeter. This model is characterized by dimensionless line tension 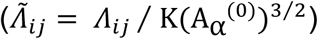 and dimensionless perimeter contractility 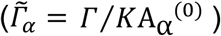 that were set respectively to 0.06 and 0.02 as in^18^. A_α_^(0)^ was initially set to 2.598 (area of an hexagon with side length of 1) for all the cells, and was subsequently modified for some cells depending on the simulated scenario (see below).

Rearrangements of the topology of the vertices (T1 transitions^36^) were allowed when two vertices i, j were located less than a minimum distance d_min_= 0.2 apart, and a movement of one of the vertices was energetically favorable such that the distance between the vertices decreases.

To generate initial conditions, we started from a set of 1801 hexagonal cells (**modelling Fig. A**), into which we introduced topological noise by allowing each cell to divide once following the Hertwig rule (i.e., by adding a new junction cleaving the cell orthogonally to its longest axis). Equation (2) was then iteratively resolved by randomly updating vertices positions 300.10^6^ times such that average cell area reached a plateau (**modelling Fig. B**). This resulted in a topology made of 3598 cells (**modelling Fig. A’**), for which both the average polygon distribution and average cell area of each polygon class qualitatively matched the ones of previous works^18,36^ (**modelling Figs. C,D**).

Next, the positions of the vertices located at the boundary of the tissue were fixed thus imposing a boundary constraint to the tissue. Two cell types were defined. In the centre of the tissue, a group of ∼150 cells was set as WT whereas all the other cells of the tissue were set as mutant (red cells, **modelling Fig. E**).

Three different scenarios of mechanical competition were explored. In the “interfacial tension” scenario (referred as Case 1, **Fig. 2A**), the line tension parameter of the junctions at the boundary between WT and mutant cells was increased by a factor λ with respect to the line tension of all the other junctions of the tissue. Four different values of λ were examined (1.0; 2.0; 5.0; 10.0). For the “differential growth” scenario (referred as Case 2, **Fig. 2A**), the resting area of the mutant cells was increased of a factor κ with respect to the resting area A^0^ of all the other cells of the tissue, with κ ε {1.0; 1.10; 1.25; 1.50}. For the third scenario (Case 3, **Fig. S4**, inset), conditions were set as in Case 1 (λ = 10.0) but with an additional condition for the junctions having one (and only one) of their vertices belonging to the boundary separating the WT and mutant populations. For these junctions, the line tension was multiplied by (−1). This allowed to simulate an increase of the adhesion at these junctions, thus reproducing our experimental observations according to which junctional tension is decreased at these junctions (**Fig. S3 G**).

Cell extrusion initiation was triggered by cell area change. We set a threshold for cell area change relative to its initial area. Four values were explored for this threshold (0.75; 0.80; 0.85, and 0.90). Once the ratio 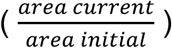 for a given cell passed below the threshold, the preferred area parameter A^(0)^ of that cell was set to 0 thus making this cell collapse further. The final step of cell extrusion was done through a T2 transition^36^: once the cell has three vertices only, and its area passes below an area threshold ( = 0.015), the three vertices were merged into a single one, and the cell was eliminated from the network.

Simulations were run for 50.10^6^ iterations (except for long runs of **Fig. S2-H,I**, 2000.10^6^ iterations), one iteration consisting in moving a randomly chosen vertex, updating its energy, and deciding to accept the movement or not. Simulation outputs were printed every 1.10^6^ iterations, thus giving outputs for 50 simulation steps.

Outputs retrieved from the simulations were: cell area change relative to initial area; circularity of the boundary between the WT cell population and the mutant population 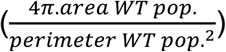; cell’s compression strength 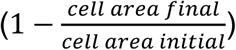; cell orientation (acute angle enclosed between the y-axis and cell’s longest axis); and proportion of cell deaths, defined as the number of cells of a given type that will die during the simulation, relative to the number of cells of that type at the beginning of the simulation. Cell type was defined according to the genotype (WT or mutant) (**modelling Fig. E**), to the distance relative to the boundary (row 1, row 2, row 3), and to the geometry of the boundary (flat or curved) (**model Fig. E’**). We defined “curved” and “flat” regions by dividing the WT patch of cells in four regions of equal size along the y-axis. The two central regions were considered as regions having a “flat” boundary, whereas the two peripheral ones were considered as having a “curved” boundary (**modelling Fig. E**). All quantification were based on 5 replicates of the simulations, obtained by changing the seed for the random walk used to update the locations of the vertices.

### Statistics

Data were not analysed blindly. No specific method was used to predetermine the number of samples. The definition of number of samples and cells considered are given in each figure legend, as well as the meaning of error bars (standard deviation or standard error of the mean (s.e.m.). p-values are calculated through group or paired t-test if the data passed normality test (Shapiro-Wilk test). Statistical tests were performed on Matlab using the ttest and ttest2 functions. For the analysis of probability of cell death as a function of distance to boundary (Fig. 1B, 3G, 4B), the paired t-test is performed by comparing averaged death probability per row with values in more distant rows from the same notum. For the analysis of death probability in “curved” vs. “flat” zone (Fig. 1L-M, 3M-P, 4H-K), the paired t-test compare death probability in curved versus flat zone within the same WT patch. Box plot are represented using the Matlab boxplot function with the central mark indicating the median, and the bottom and top edges of the box indicating the 25th and 75th percentiles, respectively.

---

**Supplementary figure 1:**
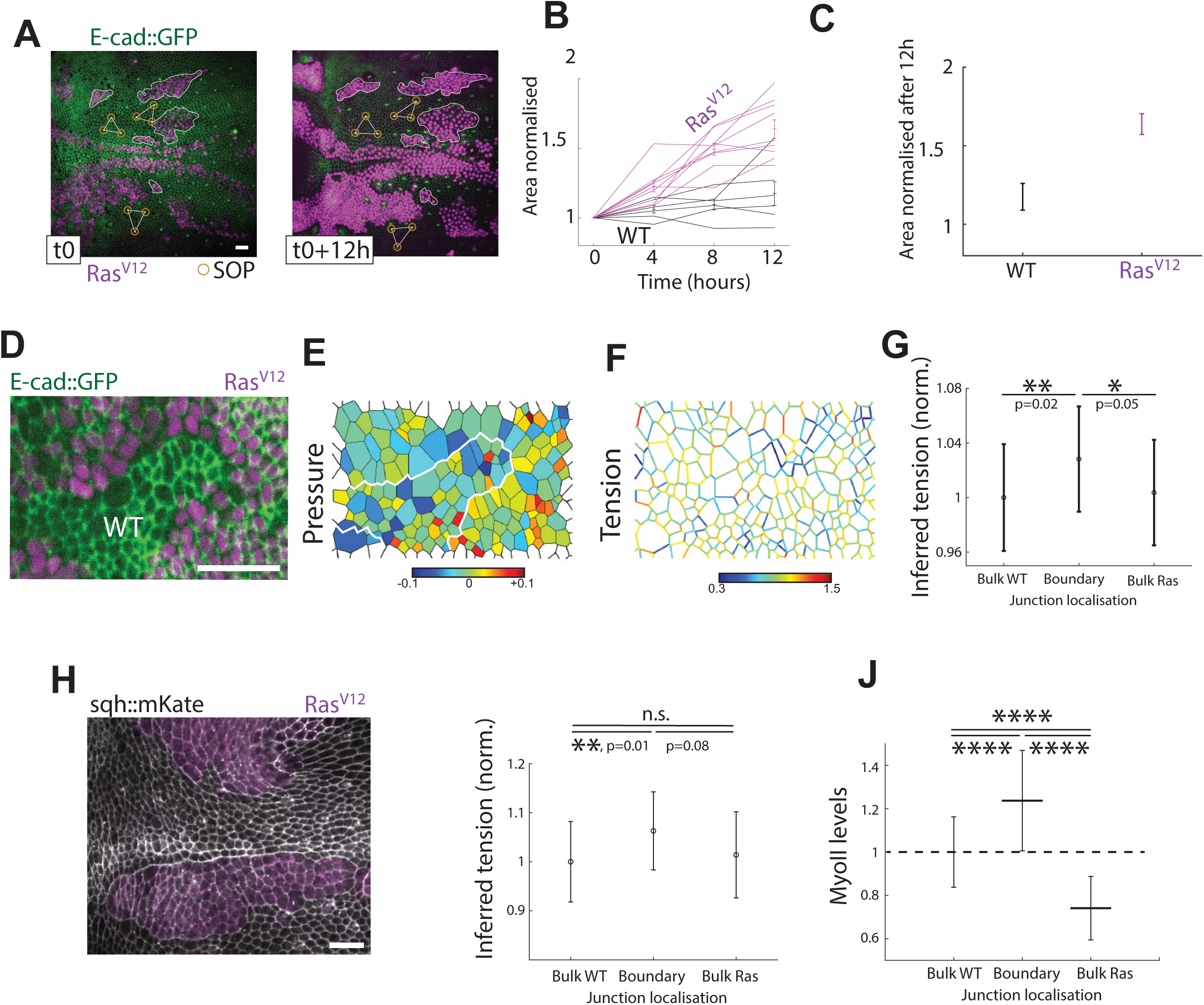
Ras^V12^ cell competition and mechanical properties (associated with Figure 1) **A-C:** Quantification of the growth of UAS-Ras^V12^ clones and WT population distant from the clones. **A:** Cells express E-cad::GFP (in green) and clones (in magenta) have been expressing UAS-Ras^V12^ for 8 hours a the onset of the movie (**A**, Left, t0) and for 12 extra hours (**A**, right). Regions used for quantification are highlighted by white areas using magenta clones or SOPs as landmarks (triangles). Scale bar is 20μm. **B:** Cell areas of group of cells (WT black, UAS-Ras^V12^ magenta) are quantified at time 0, 4, 8 and 12 hours after the beginning of the movie (t0). Data from 9 Ras clones and WT groups extracted from 2 nota. Error bars are s.e.m. **C:** Quantification of area increase after 12h (area at time t+12h over area at time t0). **D-G:** Force inference from cadherin apical junction imaging. **D:** Example image: in green E-cad::GFP; in magenta, UAS-Ras^V12^ clone. Scale bar is 20μm. **E-F:** From segmented image, we obtain maps of inferred cell pressures (**E**) and junctional tensions (**F**) (See **Star Methods**). **G:** Quantification of inferred junctional tension for junctions from 3 categories: in the bulk WT population (left), at the clone boundary in between WT and Ras^V12^ population (middle), and in the bulk of Ras^V12^ population (right). Data from 10 images extracted from 5 nota, with 267, 235 and 217 junctions respectively in each category. p values of student t-test are plotted on the graph. Error bars are s.e.m. **H-J:** MyoII distribution and force inference. **H:** Example image: in greyscale, MyoII::GFP (MRCL::GFP), in magenta, UAS-Ras^V12^ clone. Scale bar is 20μm. **I-J**: Data from 6 images extracted from 6 nota and segmented (208, 109 and 143 junctions quantified for each category). **I:** value of inferred junctional tensions obtained from force inference. p values of student t-test are plotted on the graph. Error bars are s.e.m. **J:** Averaged MyoII junctional intensity. Student t-test, **** = p<10^-4^. Error bars are s.e.m.

**Supplementary figure 2:**
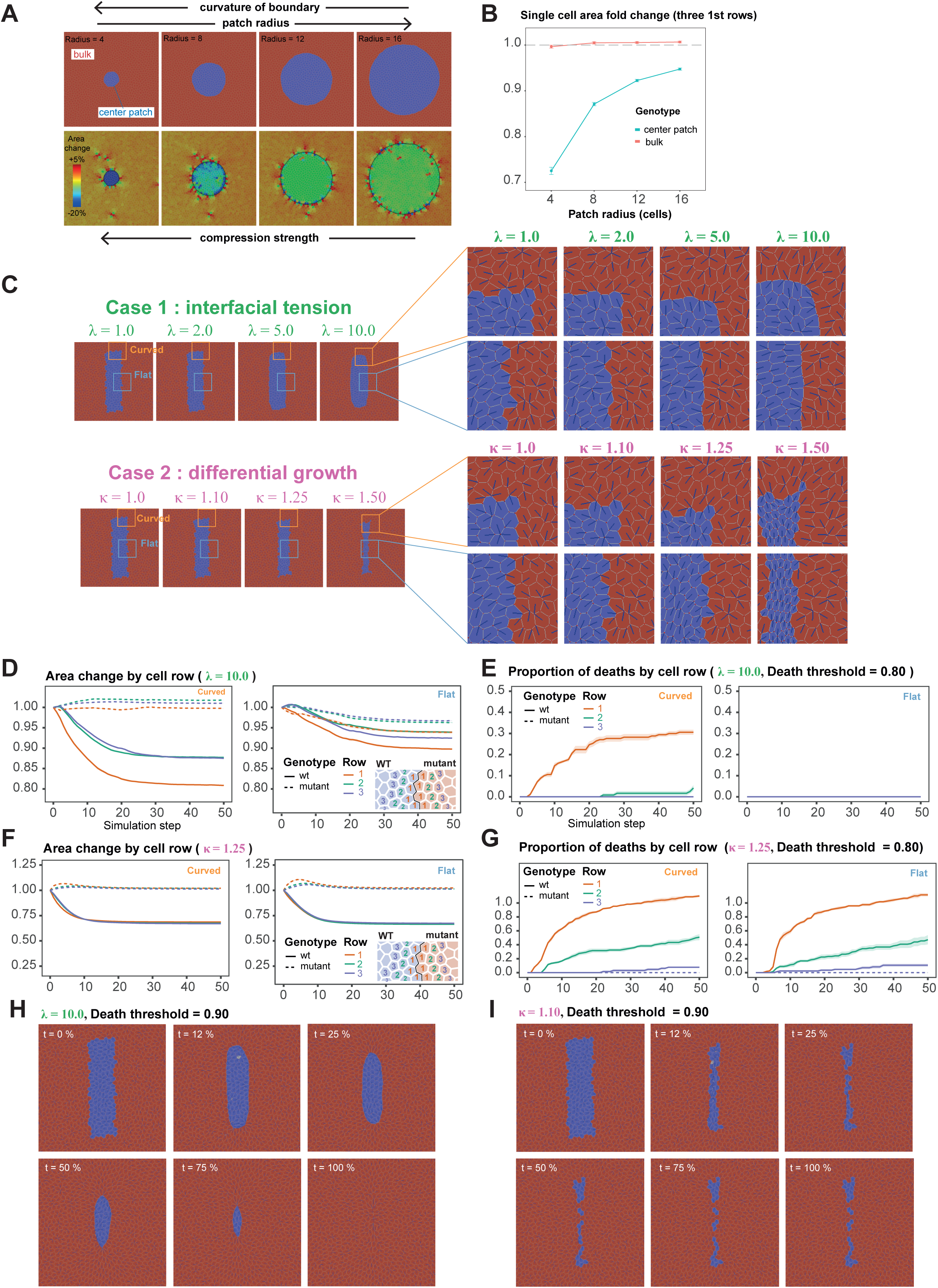
Vertex modelling of mechanical cell competition (associated with Figure 2). **A:** Illustration of the Laplace’s law in an epithelial context. Simulations were run by applying a 10-fold increase of the line tension for the junctions located at the boundary between the bulk population (orange cells) and the centre patch (blue cells). Variation in the curvature of the boundary was obtained by varying the radius of the central patch of cells, from small radius (= higher curvature) to high radius (= lower curvature). Top panels show the topology of the tissue at the final step of the simulation, and bottom panels show single cell area fold change at the final simulation step relative to area at the beginning of the simulation. Left to right: increased values of the centre patch radius (in number of cells). **B:** Quantifications from the simulations shown in A for single area fold change relative to initial area. Average +/− s.e.m. of cell area fold change is shown, as a function of cell radius and “genotype”. Areas were averaged over the three first rows of cells away from the boundary between the bulk population and the centre patch population. **C:** Close-up views of final time points from simulations shown in Figs. 2B and 2G, emphasising cell main axis orientation (blue lines) for flat (blue insets) and curved (orange insets) regions. **D:** Cell area fold change relative to initial area, as a function of simulation step, genotype, cell row, and curvature of the boundary (flat vs. curved) for a simulation of Case 1 (λ = 10.0). Cell row is given in number of rows apart from the boundary between WT and mutant populations (see diagram in the inset). **E:** Proportion of deaths as a function of simulation step, genotype, curvature of the boundary and cell row (given in number of rows apart from the boundary between WT and mutant populations, see inset in panel D). Data are shown for Case 1, λ = 10.0, death threshold = 0.80. Proportion is given relative to the number of cells of a given genotype, row and curvature region at simulation step 0. **F:** Cell area fold change relative to initial area, as a function of simulation step, genotype, cell row, and curvature of the boundary (flat vs. curved) for a simulation of Case 2 (κ = 1.25). Cell row is given in number of rows apart from the boundary between WT and mutant populations (see diagram in the inset). **G:** Proportion of deaths as a function of simulation step, genotype, curvature of the boundary and cell row (given in number of rows apart from the boundary between WT and mutant populations, see inset in panel F). Data are shown for Case 2, κ = 1.25, death threshold = 0.80. Proportion is given relative to the number of cells of a given genotype, row and curvature region at simulation step 0. **H:** Steels for six time points from a long run (2.10^9^ iterations) simulation of Case 1 (λ = 10.0, death threshold 0.90) showing complete elimination of the WT population. White cells are cells undergoing the process of T2 transition. **I:** steels for six time points from a long run (2.10^9^ iterations) simulation of Case 2 (κ = 1.10, death threshold 0.90) showing fragmentation of the WT population. White cells are cells undergoing the process of T2 transition. For D-G, quantifications are averages +/− s.e.m. obtained by running 5 times the simulations using different seeds in the Monte Carlo random walk.

**Supplementary figure 3:**
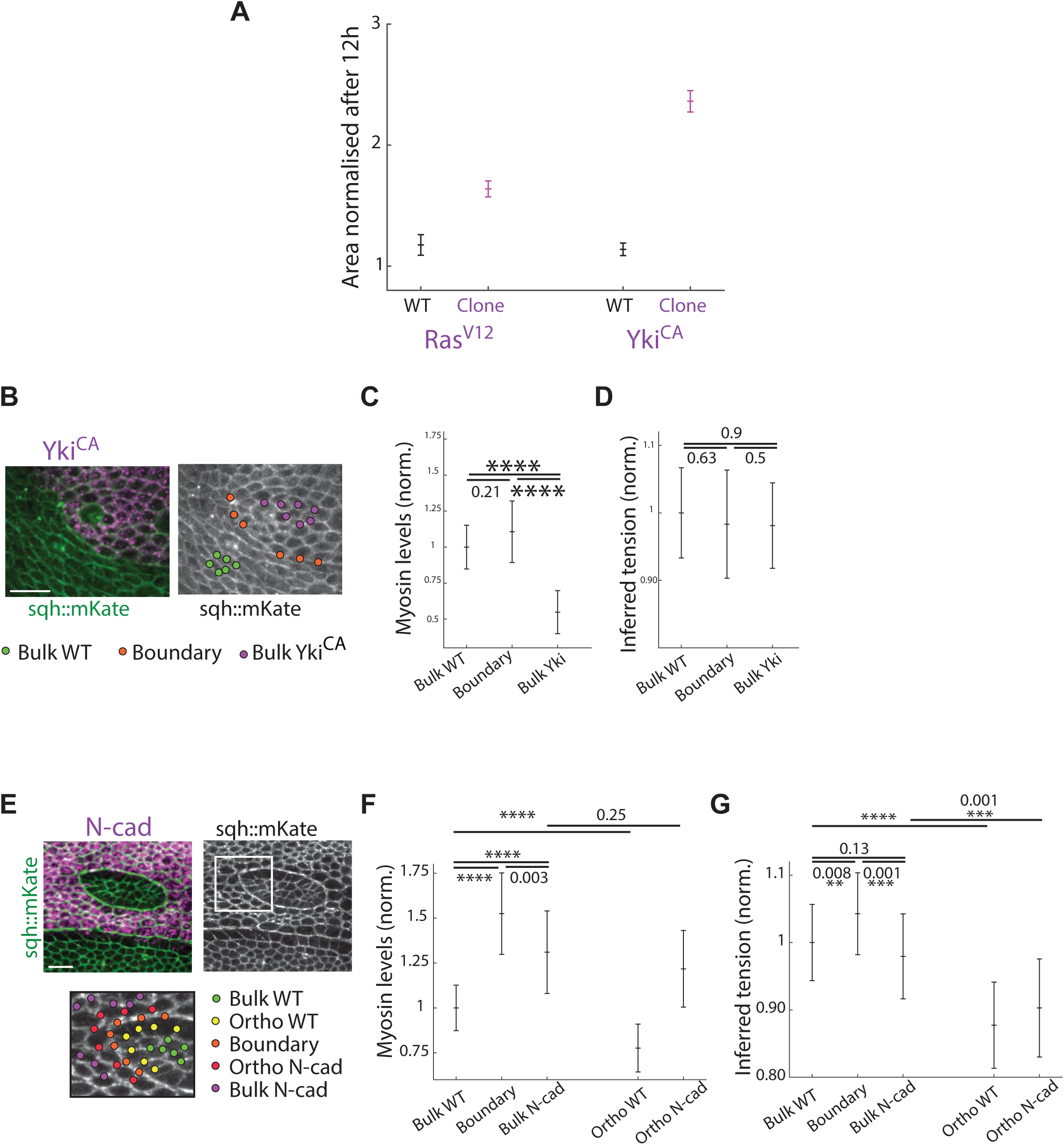
Growth and mechanical properties of Yki^CA^ clones (associated with Figure 3 and Figure 4) **A:** Cell area of group of cell, for WT cells and UAS-Ras^V12^ (left) and WT cells and UAS-Yki^CA^ (right) after 12 hours of movie (Ras^V12^, 9 groups from 2 nota; Yki^CA^ 5 groups from 2 nota). **B-D:** Force inference near Yki^CA^ clones. **B:** Example image. MyoII::GFP in green, UAS-Yki^CA^ clone in magenta. Scale bar is 20μm. **C-D**: Quantification of MyoII::GFP junctional intensity level (**C**) and inferred junctional tensions obtained from force inference (see **Star Methods**) (**D**) for junctions from 3 categories: WT bulk, clone boundaries and Yki^CA^ bulk (8 images extracted from 7 nota, junction numbers: 102, 49 and 73 respectively). Error bars are s.e.m.. Student t-test, **** = p<10^-4^. **E-G:** Force inference near UAS-N-cad clones. 2 examples of image segmented to apply the force inference pipeline. Myosin-GFP in green, N-Cad clone in magenta. Scale bar are 20μm. **F-G:** Quantification of MyoII junctional intensity (**F**) and junctional inferred tensions obtained from force inference (**G**) for junctions from 5 categories: WT bulk, clone boundary, N-cad bulk, WT orthogonal junctions touching clone boundary, N-Cad orthogonal junctions touching clone boundary, see top scheme (6 images extracted from 5 nota, junction numbers: 316, 101, 172, 97, and 66 respectively). Error bars are s.e.m.. Student t-test, **** = p<10^-4^.

**Supplementary figure 4:**
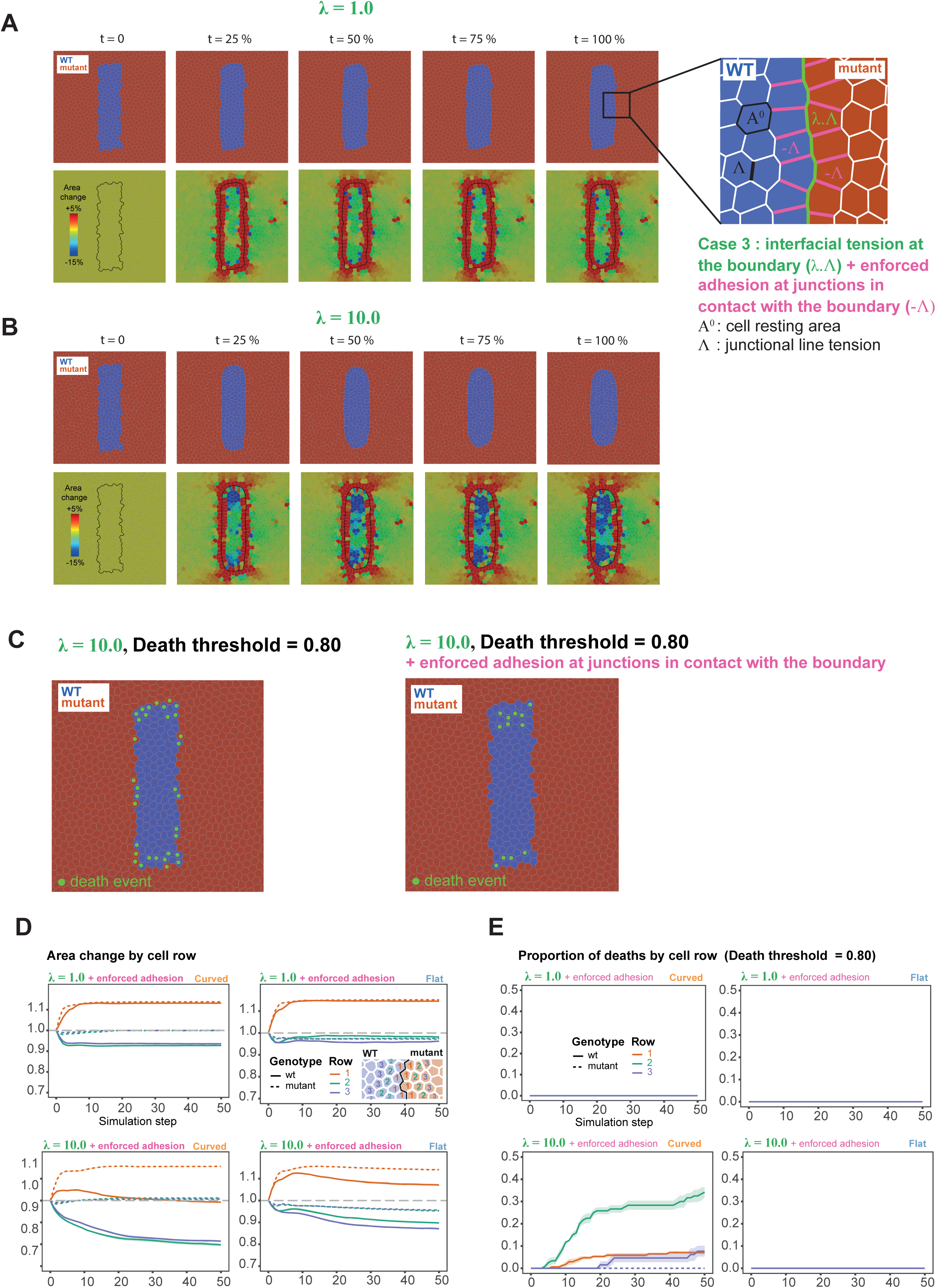
Vertex modelling of mechanical cell competition by interfacial tension increase, associated with enforced adhesion at junctions in contact with the boundary (Case 3). **A**: Steels from five time points for a control simulation of Case 1 (λ = 1.0) and with the addition of enforced adhesion at the junctions in contact with the boundary. Top panels show the topology of the tissue over simulation steps (from initial (0 %, left) to final (100 %, right)), and bottom panels show single cell area fold change relative to area at the beginning of the simulation. Inset: to simulate our assumption of enforced adhesion resulting in the lengthening of junctions observed in the vicinity of the boundary of UAS-N-cad clones, we set a specific line tension condition for the junctions in contact with the boundary (magenta junctions in the inset). These junctions were assigned a negative value of Λ, thus making the lengthening of the junction mechanically favourable. Running a control simulation of Case 1 (λ = 1.0) with this additional condition was sufficient to lengthen the junctions in contact of the boundary, thus increasing cell area of cells in the contact with the boundary (bottom panels). Interestingly, this addition was also sufficient to slightly smooth the boundary between WT and mutant cells, as well as to generate slight compaction within the WT and mutant populations away from the cell rows in direct contact with the boundary. **B:** Steels from five time points of a simulation of mechanical cell competition by interfacial tension (Case 1, λ = 10.0) and with the addition of enforced adhesion at the junctions in contact with the boundary. Top panels show the topology of the tissue over simulation steps, and bottom panels show single cell area fold change relative to area at the beginning of the simulation. **C:** Back tracking of dying cells. Cells that will die during the simulation are highlighted on the topology at simulation step 0 (green dots). Left panel shows the pattern of dying cells for Case 1, λ = 10.0, and right panel shows the pattern for Case 1, λ = 10.0 + enforced adhesion at junctions in contact with the boundary (Case 3). Note the quasi absence of dying cells for cells in direct contact with the boundary in Case 3 (right) compared to Case 1 (left). **D:** Cell area fold change relative to initial area, as a function of simulation step, genotype, cell row, curvature of the boundary (curved, left panels vs. flat, right panels) and λ (1.0, top panels vs. 10.0, bottom panels) in simulations with enforced adhesion at junctions in contact with the boundary (Case 3). Cell row is given in number of rows apart from the boundary between WT and mutant populations (see diagram in the inset). **E:** Proportion of deaths as a function of simulation step, genotype, cell row, curvature of the boundary (curved, left panels vs. flat, right panels) and λ (1.0, top panels vs. 10.0, bottom panels) in simulations with enforced adhesion at junctions in contact with the boundary (Case 3) and death threshold set to 0.80. Cell row is given in number of rows apart from the boundary between WT and mutant populations (see diagram in the inset of D). Proportion is given relative to the number of cells of a given genotype, row and curvature region at simulation step 0. For D-E, quantifications are averages +/− s.e.m. obtained by running 5 times the simulations using different seeds in the Monte Carlo random walk.

**Supplementary figure 5:**
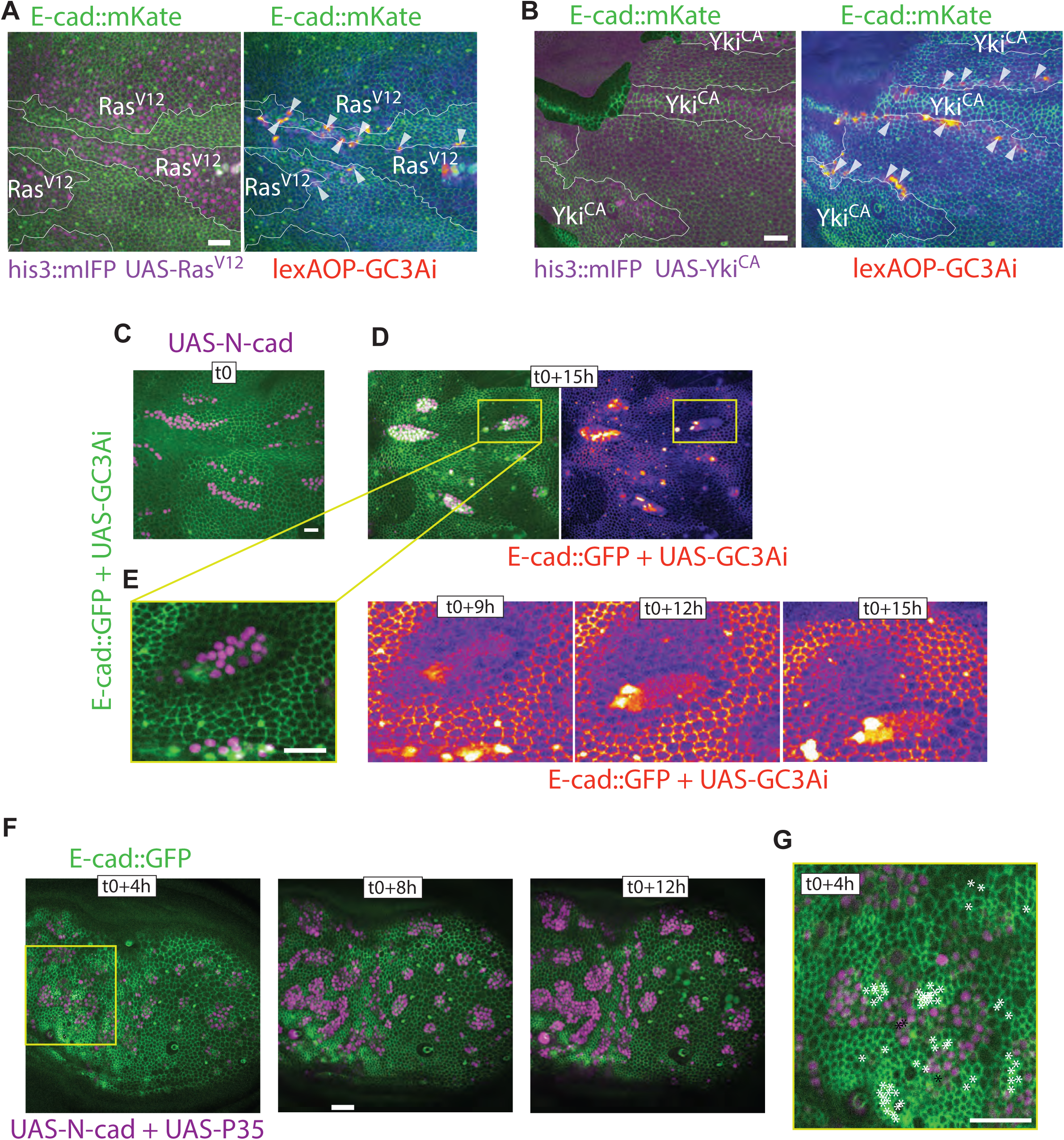
Caspase activity during UAS-Ras^V12^, UAS-Yki^CA^ and UAS-N-cad mechanical cell competition (associated with Figure 1, 3 and 4). **A:** Representative snapshot (local z-projection) of movie showing E-cad::mKatex3 in green, and Ras^V12^ clones in magenta (left) and the caspase sensor lexAOP-GC3Ai in “fire red”(right, ubiquitous expression driven by act-lexA). Some cells with high caspase activity are highlighted with white arrowheads. Scale Bar is 20 μm. **B:** Representative snapshot (local z-projection) of movie showing E-cad::mKatex3 in green, and Yki^CA^ clones in magenta (left) and the caspase sensor lexAOP-GC3Ai in “fire red”(right, ubiquitous expression driven by act-lexA). Some cells with high caspase activity are highlighted with white arrowheads. Scale Bar is 20 μm. **C-E:** Representative example of movies showing E-cad::GFP and UAS-GC3Ai in green or “fire red”, UAS-N-cad clones in magenta before (**C**) and after 15 hours (**D**). Of note, here GC3Ai caspase sensor is only expressed in the clones (contrary to **A** and **B**). **E:** Zoom on one group of N-cad cells, and snapshots of E-cad::GFP and UAS-GC3Ai every 3 hours. Scale Bar is 20 μm. **F:** Representative example of a movie showing E-cad::GFP in green and UAS-N-Cad + UAS-p35 (effector caspase inhibitor) clones in magenta for 3 time points separated by 4 hours. Most extrusions appearing in the yellow square during the whole movie are back tracked and represented on timepoint t0+4h as asterisks (*) and in its zoom (G). White stars are dying WT cells, black stars UAS-N-Cad, UAS-p35 dying cells. Note that there is hardly any extruding cell in the N-cad, UAS-p35 clone. Representative of 2 movies. Scale bar is 20μm.

**Supplementary figure 6:**
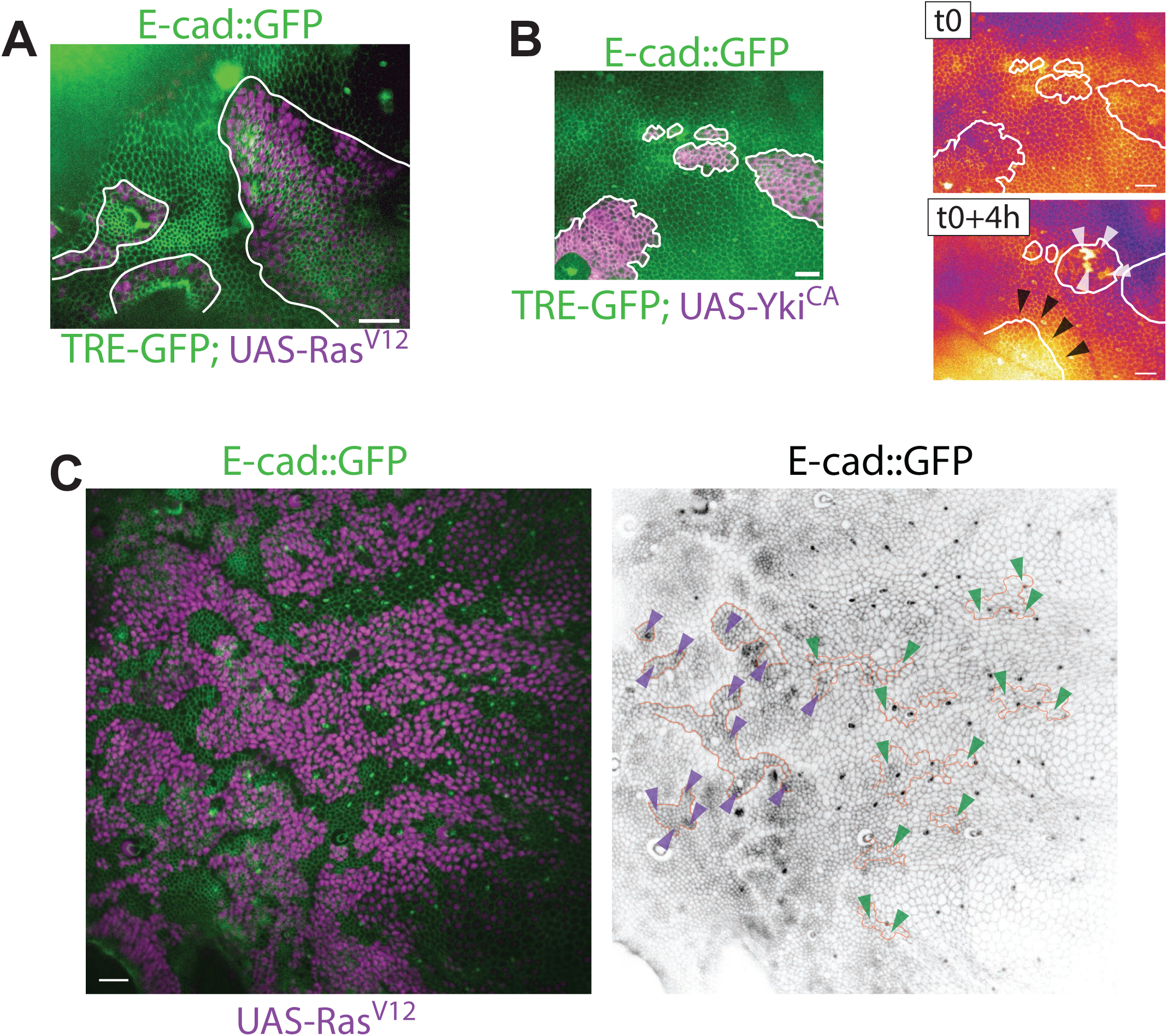
JNK activity during Ras^V12^ and Yki^CA^ mechanical competition and tissue context-dependent sorting (associated with Figure 1 and Figure 3). **A-B:** JNK activity during mechanical competition mediated by Ras^V12^ and Yki^CA^ is not specifically localised at compacted and/or dying cells. E-cad::GFP and TRE-GFP (transcriptional reporter of JNK activity) in green and clones in magenta. **A:** Representative example of 4 experiments for which pupae are imaged 20h after induction of Ras^V12^ in clones where no clear JNK activation was observed (even in compacted cells) **B:** Representative example of a movie in the case of Yki^CA^ clones. Left: Snapshots of E-cad::GFP and TRE-GFP (green, Yki^CA^ cells in magenta), right panels : E-cad::GFP and TRE-GFP shown in “fire red” for 2 timepoints separated by 4 hours. Black arrowheads show a clone boundary. White arrowheads show few sporadic cells activating JNK in one clone. Scale bars are 20μm. **C:** The degree of Ras^V12^ clone sorting is region specific in the pupal notum. Representative example of E-cad::GFP in green (left) or grayscales (right) and UAS-Ras^V12^ clones in magenta. Posterior, left, Anterior, right. Scale bar is 20μm. Right, highlight of contours of some group of WT cells. Magenta arrowheads: WT patches with clear sorting behaviour (mostly in the posterior region). Green arrowheads: WT group with no obvious sorting behaviour (mostly anterior).

## Methods Modelling – figure

**Modelling figure.**
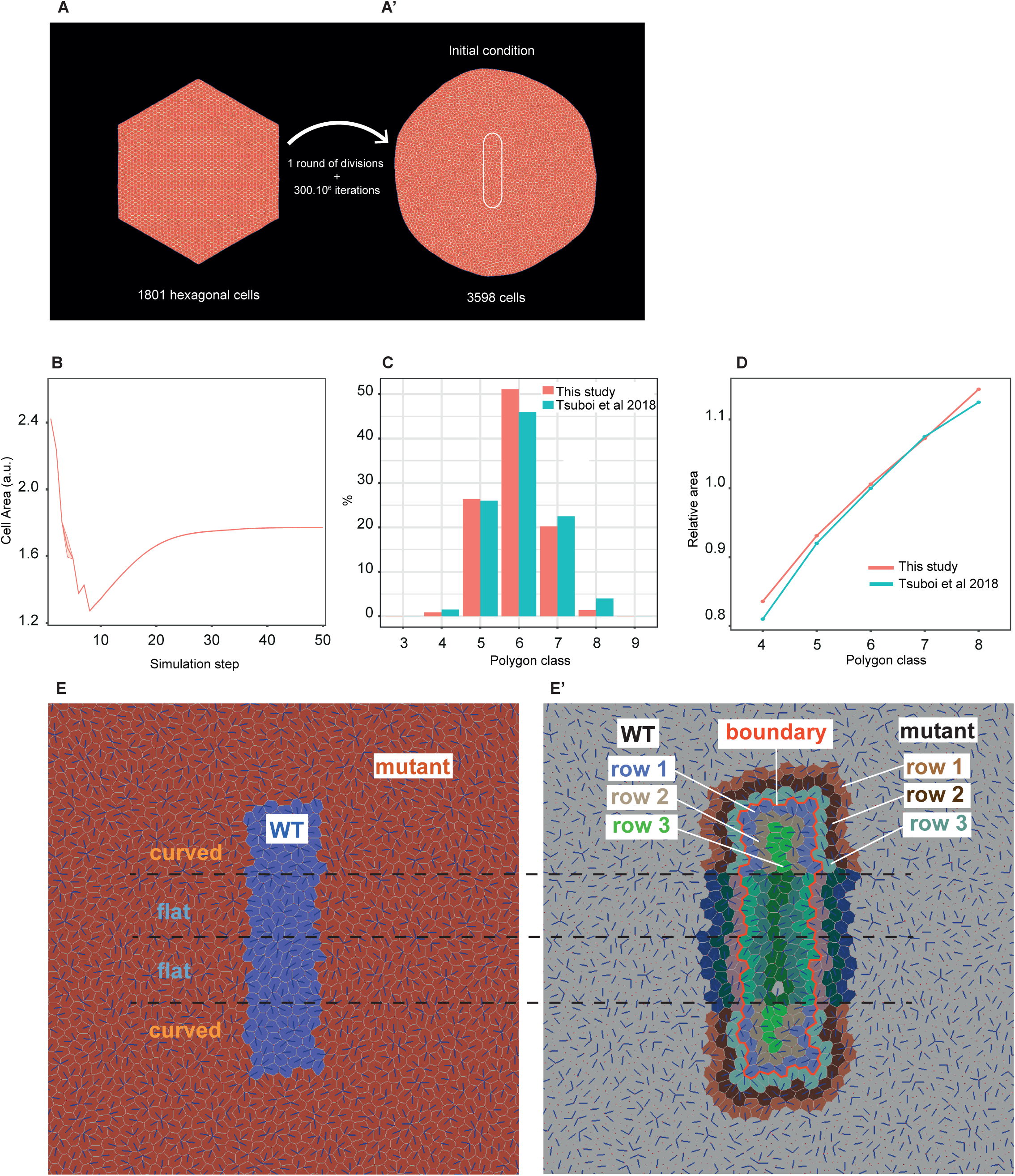
Description of the parameters of the vertex model. **A-A’:** Generation of initial conditions for the simulations. The white ellipse in A’ denotes the area of the WT patch of cells. **B:** Evolution of the average cell area during the 300.10^6^ iterations of the simulation ran to generate the initial condition topology (A-A’). The average area reaches a plateau indicating that the networks is close to a steady-state condition. **C:** Frequency distribution of polygon classes (i.e. number of junctions per cell) found in our initial condition topology, compared to the one in^18^. **D:** Average area of cells for each polygon class normalised by the average cell area of the total cell population in our initial condition topology (=Relative area), as a function of polygon class (i.e. number of junctions per cell). **E-E’**: Steel from the initial condition topology showing the definition of the cell types. Cell type was defined according to cells’ genotype (WT or mutant) (E), the curvature (“curved” or “flat”) (E); and number of cell rows apart from the boundary between the WT and the mutant cells populations (E’).

## Supplementary movie legends

**Movie S1:** Movie related to **Fig. 1A**. Notum local z-projection with UAS-Ras^V12^ clones imaged over 22 hours in which dying cells localisation is highlighted by white circles (from 40 minutes before to 40 minutes after death). In green, E-cad::GFP; In magenta, nuclei of clonal cells overexpressing Ras^V12^. Scale bar is 20μm.

**Movie S2:** Movie related to **Fig. 1G**. 2 examples (left and right) of WT populations (in the centre) being compacted and eliminated by surrounding cells overexpressing Ras^V12^. In green, E-cad::GFP; In magenta, nuclei of clonal cells overexpressing Ras^V12^. Scale bar is 20μm.

**Movie S3:** Movie related to **Fig. 1J**. Example of WT cells (in the centre) being compacted and eliminated by surrounding cells overexpressing Ras^V12^ in which dying cells localisation is highlighted by white circles (from 40 minutes before till death). In green, E-cad::GFP; In magenta, nuclei of clonal cells overexpressing Ras^V12^. Scale bar is 20μm.

**Movie S4:** Movie related to **Fig. 2B**. Vertex simulation of cell deformation induced by an increase of line tension at clone boundary (blue cells). Blue lines are aligned with the main axis of cell elongation. From left to right: λ=1.0, 2.0, 5.0 and 10.0. The bottom movies show the relative change of cell area (red: cell expansion, blue: cell compaction), the black line shows the clone contour.

**Movie S5:** Movie related to **Fig. 2G**. Vertex simulation of cell deformation induced by neighbouring cell growth driven by a progressive increase of their resting area (red cells). Blue lines are aligned with the main axis of cell elongation. From left to right: κ=1.0, 1.1, 1.25 and 1.5. The bottom movies show the relative change of cell area (red: cell expansion, blue: cell compaction), the black line shows the clone contour.

**Movie S6:** Movie related to **Fig. 2E,F,J,K**. Vertex simulation of cell deformation and cell elimination (white cell) induced by an increase of line tension at clone boundary (top, λ=10.0) or upon increase of the resting area of neighbouring cells (red cell, bottom, κ=1.25). From left to right, cell death compaction threshold of 0.75, 0.8, 0.85 and 0.9.

**Movie S7:** Movie related to **Fig. 3E**. Notum local z projection with UAS-Yki^CA^ clones imaged over 19 hours in which dying cells localisation is highlighted by white circles (from 40 minutes before to 40 minutes after death). In green, E-cad::GFP; In magenta, nuclei of clonal cells overexpressing Yki^CA^. Scale bar is 20μm.

**Movie S8:** Movie related to **Fig. 3H**. 2 examples (left and right) of WT populations (in the centre) being compacted and eliminated by surrounding cells overexpressing Yki^CA^. In green, E-cad::GFP; In magenta, nuclei of clonal cells overexpressing Yki^CA^. Scale bar is 20μm.

**Movie S9**: Movie related to **Fig. 3M**. Example of WT cells (in the centre) being compacted and eliminated by surrounding cells overexpressing Yki^CA^ in which dying cells localisation is highlighted by white circles (from 40 minutes before till death). In green, E-cad::GFP; In magenta, nuclei of clonal cells overexpressing Yki^CA^. Scale bar is 20μm.

**Movie S10:** Movie related to **Fig. 4C**. Notum local z projection with UAS-N-cad clones imaged over 23 hours in which dying cells localisation is highlighted by white circles (from 40 minutes before to 40 minutes after death). In green, E-cad::GFP; In magenta, nuclei of clonal cells overexpressing N-Cad. Scale bar is 20μm.

**Movie S11:** Movie related to **Fig. 4G**. 2 examples (left and right) of WT populations (right) and UAS-N-cad clone (left) being surrounded by the other genotypes in which dying cells localisation is highlighted by white circles (from 40 minutes before till death). In green, E-cad::GFP; In magenta, nuclei of clonal cells overexpressing N-cad. Scale bar is 20μm.

**Movie S12:** Movie related to **Fig. S4**. Vertex simulation of cell deformation induced by an increase of line tension at clone boundary (blue cells) and upon enforced at adhesion in orthogonal junction (hence leading to junction length increase). Blue lines are aligned with the main axis of cell elongation. Left: λ=1.0, Right, λ=10.0. The bottom movies show the relative change of cell area (red: cell expansion, blue: cell compaction), the black line shows the clone contour.

**Movie S13:** Movie related to **Fig. S5**. Visualisation of caspase activity using the GC3Ai sensor during mechanical cell competition over 10 hours. From Top to bottom: Ras^V12^ clones (top), Yki^CA^ clones (middle) and N-cad clones (bottom). (Left) In green, E-cad::GFP; In magenta, nuclei of clonal Gal4 expressing cells. (Right) In green, E-cad::GFP; In “fire-red” LUT, tub-lexA, lexAOP-GC3Ai (top and middle, ubiquitous expression) and UAS-GC3Ai (bottom, expression in UAS-N-cad clone only) signals. Scale bar is 20μm.

## Notes

### Competing Interest Statement

The authors have declared no competing interest.

